# EBAX-1/ZSWIM8 destabilizes miRNAs resulting in transgenerational memory of a predatory trait

**DOI:** 10.1101/2024.09.10.612280

**Authors:** Shiela Pearl Quiobe, Ata Kalirad, Waltraud Röseler, Hanh Witte, Yinan Wang, Christian Rödelsperger, Ralf J. Sommer

**Affiliations:** Department for Integrative Evolutionary Biology, Max Planck Institute for Biology Tübingen; Max-Planck Ring 9, Tübingen, 72076, Germany

## Abstract

Environmental influences on traits and associated transgenerational epigenetic inheritance have widespread implications, but remain controversial and underlying mechanisms poorly understood. We introduce long-term environmental induction experiments on alternative diets in *Pristionchus pacificus*, a nematode exhibiting mouth-form plasticity including predation, by propagating 110 isogenic lines for 101 generations with associated food-reversal experiments. We found dietary induction and subsequent transgenerational memory of the predatory morph and identified a role of ubiquitin ligase EBAX-1/ZSWIM8 in this process. *Ppa-ebax-1* mutants have no memory and *Ppa-*EBAX-1 destabilizes the clustered microRNA family *miR-2235a/miR-35.* Deletions of a cluster of 44 identical *miR-2235a* copies resulted in precocious and extended transgenerational inheritance of the predatory morph. These findings indicate that EBAX-1/ZSWIM8 destabilizes miRNAs resulting in transgenerational memory, suggesting a role for target-directed miRNA degradation.

## Main Text

Environmental influences on trait inheritance in animals including humans have long been suggested but remain controversial, largely because of the many confounding effects encountered when measuring such phenotypes over multiple generations (*1*, *2*). However, laboratory studies in model organisms, most notably the nematode *Caenorhabditis elegans* with its short generation time, isogenic propagation and simple husbandry, have provided powerful demonstrations of transgenerational epigenetic inheritance (TEI) and started to provide mechanistic insight (for review see (*3–6*)). Indeed, a growing body of evidence suggests that the environment can induce various phenotypic changes that are transmitted across generations. In this context, intergenerational and transgenerational inheritance are distinguished, with the former describing transmission that lasts for only two, whereas the latter describes memory that lasts for at least three generations beyond parental induction (*3*). Already transient, short-term environmental variation can induce TEI in *C. elegans* (*7–14*). For example, a single exposure to elevated temperature, starvation or pathogens induces changes of small interfering (si)RNAs that last for several generations (*12*, *15–19*). Also, Argonaute proteins and histone modifications were shown to play a crucial role in siRNA signaling in *C. elegans* (*7–9*). However, a sophisticated machinery typically helps resetting such nongenetic effects after 3-5 generations (*20*).

Nonetheless, several experimental challenges impede the study of TEI and its long-term significance for the organism (*6*, *21–23*). We try to overcome some of these challenges through two innovations in experimental design. First, we assay a natural, organismal readout in form of developmental plasticity in nematode feeding structures, which results in a binary morphological phenotype among genetically identical organisms (Fig. 1A). Second, we use a study system borrowed from experimental evolution (*24*) for the analysis of environmental influence on plastic trait formation and potential TEI. Specifically, we study the second nematode model organism *Pristionchus pacificus* that shares with *C. elegans* its easy husbandry, short generation time and hermaphroditic mode of reproduction resulting in isogenic cultures (*25*, *26*). However, unlike *C. elegans*, *P. pacificus* exhibits a mouth-form dimorphism that is an exemplar of developmental plasticity with morphological and behavioral implications (Fig.1A) (*27–29*). By definition, the environment influences which of the two alternative phenotypes is expressed in a given individual as part of the ‘natural’ biology of this organism, providing an ideal approach to investigate potential systems of TEI.

**Fig. 1.**
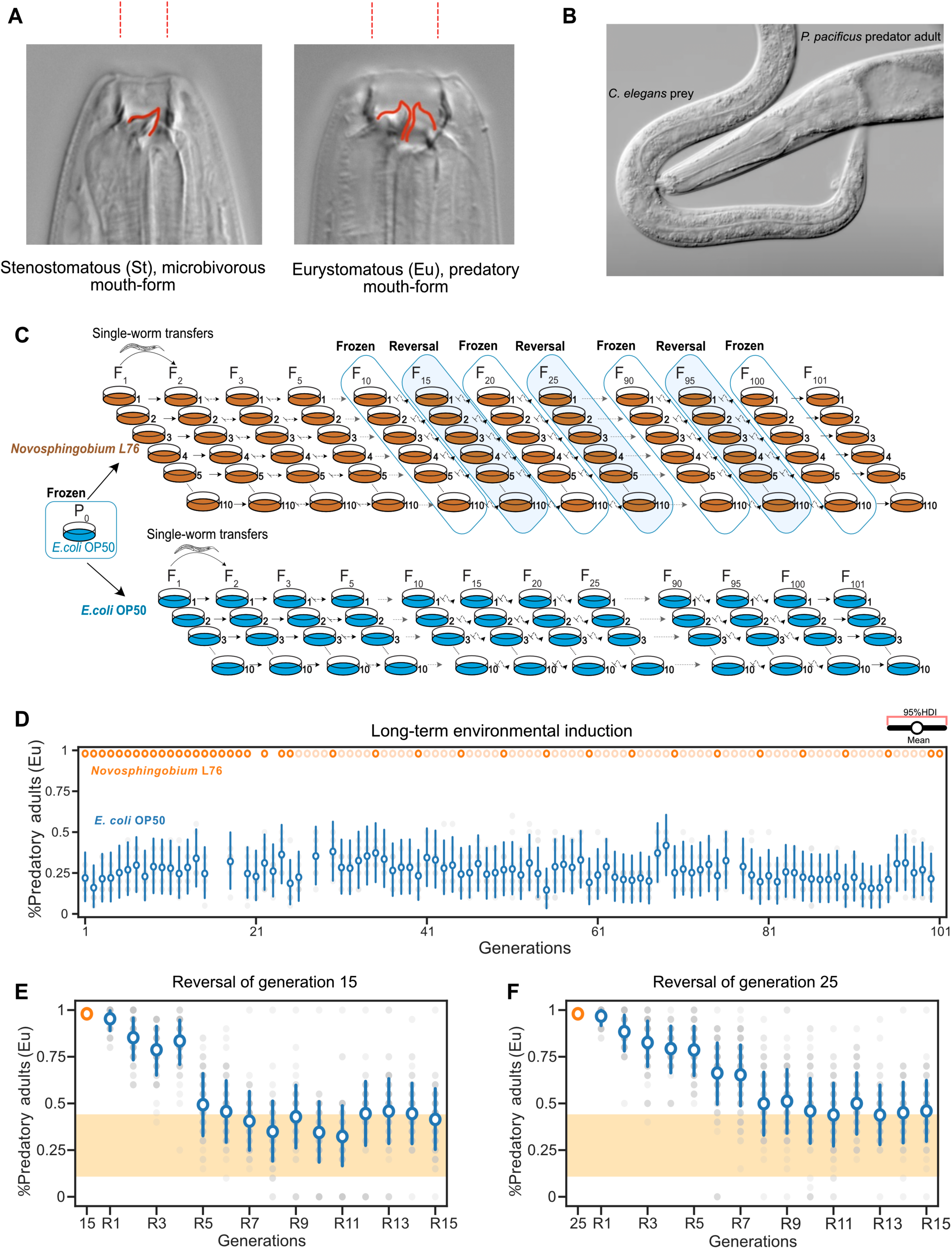
Long-term environmental induction approach on alternative diets shows induction and transgenerational epigenetic inheritance of the predatory morph in *Pristionchus pacificus*. (**A**) Mouth dimorphism of *P. pacificus*. The eurystomatous (Eu) morph has a wide mouth with two claw-like teeth and feeds on both bacteria and other nematodes, whereas the stenostomatous (St) morph has a narrow mouth with a single flint-like dorsal tooth and feeds only on microbes. (**B**) Adult Eu *P. pacificus* devouring a *C. elegans* larval prey. (**C**) Experimental design for long-term environmental induction using distinct diets. 110 parallel lines derived from a single hermaphrodite were exposed to a *Novosphingobium* diet with 10 control lines remaining on the standard *E. coli* food source. All lines were propagated for 101 generations by single-worm descent. Periodic food reversal experiments back to the standard *E. coli* diet were performed all 10 generations starting in generation F15. In addition, lines were frozen all 10 generations starting in generation F10. (**D**) Mean probability of the predatory mouth-form on *Novosphingobium* (orange) and *E. coli* (blue) during the long-term environmental induction experiment. Like in all following graphs, the y-axis indicates the observed Eu mouth form frequency and x-axis shows the number of generations. The 95% highest density interval (HDI) of the mean for every generation was used to visualize the probability of observing the Eu morph. (**E**) Mean probability of predatory mouth-form on *E. coli* after 15 generations of exposure to *Novosphingobium*. (**F**) Mean probability of predatory mouth-form on *E. coli* after 25 generations of exposure to *Novosphingobium*. The 95% HDI for the means was estimated using a Bayesian hierarchical model and indicates the probability of expressing the Eu morph inferred from the experimental data. The yellow region displays the upper and lower limit of the baseline Eu response of RSC011 (0.115 ≤HDI(^θ̅^_Control_)≤ 0.436), averaged across 101 generations on *E. coli* (see Materials and Methods for details).

During the last decade, *P. pacificus* mouth-form plasticity has emerged as one of the study systems in plasticity research (*30*). During postembryonic development, genetically identical individuals adopt either the narrow-mouthed ‘stenostomatous’ (St) morph with a single dorsal tooth, or the wide-mouthed ‘eurystomatous’ (Eu) form with a claw-like dorsal and an opposing sub-ventral tooth (Fig. 1A). Importantly, Eu animals are omnivores that can feed on bacteria, fungi and other nematodes as predators, whereas St worms are strict bacterial feeders (Fig. 1B) (*31*). Thus, the mouth-form decision of an individual animal has important behavioral and ecological consequences (*32*). This instance of plasticity, where environmental stimuli affect the likelihood of an individual developing one of the two alternative states, can be characterized as a bi-stable developmental switch (*33*, *34*), distinct from other cases of more-or-less deterministic phenotypic plasticity (e.g., wing polyphenism in butterflies (*35*)). The gene regulatory network (GRN) regulating mouth-form plasticity in *P. pacificus* has been characterized in detail and involves a complex mixture of highly conserved as well as rapidly evolving genes (*28*, *36–44*). In addition, multiple environmental cues influence mouth-form development. For example, culture methods (*45*), temperature (*46*) and small-molecule metabolites (*47*) were shown to influence the Eu *vs.* St mouth-form ratio of isogenic *P. pacificus* strains. More recently, changes in diet were also found to profoundly affect the mouth form, establishing an easy study system for experimental manipulation (*48*). Here, we used long-term environmental induction through distinct diets by propagating 110 *P. pacificus* isogenic lines for 101 generations with associated food-reversal experiments.

### A long-term environmental induction experiment to visualize plasticity of feeding structures

We borrowed the concept of experimental evolution (*24*) to study the influence of diet on *P. pacificus* mouth-form plasticity. Specifically, we established a long-term environmental induction experiment for 101 generations with regular food reversals, to investigate the consequences of dietary change (Fig. 1C). We previously showed that multiple bacterial isolates from nematode-associated environments can alter *P. pacificus* mouth-form ratios and predatory behaviors (*48*, *49*). For example, growing *P. pacificus* for one generation on *Novosphingobium* L76 increased killing efficiency relative to *Escherichia coli*-grown worms and also changed mouth-form ratios towards Eu in certain *P. pacificus* strains (*48*, *49*). Therefore, we performed the long-term environmental induction experiment by switching naïve *P. pacificus* cultures that were always grown on the standard laboratory food source *E. coli* OP50 to *Novosphingobium* (Fig. 1C). We used the strain *P. pacificus* RSC011 from La Réunion Island for the experiment because this strain is preferentially non-predatory (30%Eu:70%St) when grown on *E. coli* (*48*).

*P. pacificus* RSC011 is isogenic after 10 generations of original inbreeding and was subsequently frozen as a reference strain. However, this strain has not undergone an inordinately long culture period in the laboratory that could have resulted in domestication. Together, the long-term environmental induction setup relies on i) the propagation of isogenic cultures by self-fertilization, ii) simple and rapid worm husbandry on monoxenic bacterial diets, iii) the formation of large brood sizes for phenotyping, and iv) the natural morphological readout of feeding structure plasticity.

To start the experiment, we singled out the complete brood of a single RSC011 adult hermaphrodite derived from an *E. coli* OP50 culture (Fig. 1C). We established 110 parallel lines on the *Novosphingobium* diet and 10 control lines that remained on standard *E. coli* food. All lines were propagated for a total of 101 generations. In each generation, a single healthy worm at the J4 juvenile stage was propagated to a new plate with the respective diet (Fig. 1C). We used single worm descend to maximize founder effects and the chance that rare and stochastic, epigenetic events can manifest themselves in a population. Indeed, recent studies in *C. elegans* revealed that animals can stochastically assume transgenerational epigenetic states (*13*, *50*). This approach is by design different from selection line experiments, where several animals are propagated together in order to draw on standing genetic variation already present in a population. In contrast, in our experiment all animals are genetically identical and propagation by single worm descend allows capturing stochastic epigenetic variation, although the accumulation of genetic mutations cannot be ruled out.

The naïve RSC011 culture, from which the single worm for starting the experiment was taken, was frozen as reference. Similarly, all 110 lines were frozen all 10 generations (F10, F20 etc.). Genome sequencing of some of these frozen lines after an exposure to *Novosphingobium* for 100 generations revealed a mutation rate of 1-3 x 10^-9^ (fig. S1K), which is similar to the previously reported mutation frequency in *P. pacificus* (*51*). Given the genome size of *P. pacificus,* this results in roughly one mutation in 10 generations of the experimental setup. With a generation time of four days, the complete long-term environmental induction experiment was carried out in approximately one year.

### *Novosphingobium* induces the predatory mouth form and causes massive changes in gene expression

The 110 sister lines of *P. pacificus* RSC011 developed a mouth-form ratio of 100% Eu animals already in the first generation cultured on *Novosphingobium* (Fig. 1D). In contrast, the probability of developing the Eu mouth form (θ) in *E. coli* control lines remained low in this generation and throughout the entire experiment (0.115 ≤HDI(θ)≤ 0.436) (Fig. 1D). The immediate change in mouth-form ratio on *Novosphingobium* remained stable in all lines and in all subsequent generations with an all-Eu phenotype (Fig.1D). In comparison to *E. coli*-grown worms, dietary induction of the predatory mouth form is immediate (starting in generation 1), systemic (occurred in all 110 lines) and complete (100% Eu). These findings are further supported by transcriptomic studies upon *Novosphingobium* exposure (fig. S2A; Data S2, A to D). Specifically, RNA-seq analysis identified 3,043 differentially expressed genes (adjusted P-value <0.01) between both bacterial diets (P0 on *E. coli vs.* F3 on *Novosphingobium*). These genes are mostly enriched in Collagens, Cadherins and Vitellogenin (fig.S2A; Data S2A). Thus, exposure to *Novosphingobium* induces a massive transcriptomic response in RSC011 animals.

### Dietary induction of the predatory morph shows transgenerational epigenetic inheritance

Dietary effects can cause TEI in nematodes (*12*, *17*, *52*). Therefore, we started to revert all 110 lines back to standard *E. coli* OP50 food in generation F15 (Fig. 1C and Fig. 1E). For that, we transferred one J4 juvenile per line to an NGM agar plate supplemented with kanamycin and used a kanamycin-resistant *E. coli* strain for one generation. Kanamycin-supplemented plates and the kanamycin-resistant *E. coli* strain, which is a direct derivative of the OP50 strain, on their own had no effect on *P. pacificus* mouth form (fig. S1J). Subsequently, all lines were cultured for 14 additional generations on *E. coli* OP50 (Fig.1E). We repeated these experiments all 10 generations with a total of nine reversals between F15 and F95 (Fig.1, E and F and fig. S1, C to I).

In the F15 reversal experiment, we observed high probability of developing the Eu mouth form in the first generation (0.89 ≤HDI(θ_F15R1_)≤ 0.953) (F15R1), and in the three subsequent generations (0.733 ≤HDI(θ_F15R2_)≤ 0.957, 0.652 ≤HDI(θ_F15R3_)≤ 0.913, and 0.709 ≤HDI(θ_F15R4_)≤ 0.946) (Fig.1E). Only in generation F15R5, the mean mouth-form ratio dropped below 50% Eu (θ̄_F15R5_ =0.493, 0.327 ≤HDI(θ_F15R5_)≤ 0.66), which is not credibly different from the baseline level of naïve worms (0.115 ≤HDI(θ_Control_)≤ 0.436) (see Materials & Methods for details). These findings indicate a TEI of the Eu mouth form that lasts for four generations.

When we repeated the reversal experiment in generation F25, we found even stronger TEI that also lasted for more generations. Specifically, in F25R1 all but one line had at least 85% Eu animals (θ̄_F15R1_ =0.968, 0.917 ≤HDI(θ_F25R1_)≤ 0.999) (Fig.1F). The probability of developing the Eu mouth form remained high in the F25R5 generation (0.649 ≤HDI(θ_F25R5_)≤ 0.913) and only decreased considerably in generation F25R8 (0.33 ≤HDI(θ_F25R8_)≤ 0.667), whereas such a value was observed already in F15R5 (0.327 ≤HDI(θ_F15R5_)≤ 0.66) (Fig.1, E and F). Thus, comparing the F15 and F25 reversal experiments suggests extended TEI of the predatory mouth form correlating with the length of dietary induction. Analysis of the subsequent reversal experiments indicated strong TEI and the mean proportion of Eu animals at generation 15 after the reversal to fluctuate around 50% -60% (e.g., 0.349 ≤HDI(θ_F15R35_)≤ 0.664, ≤0.32≤HDI(θ_F15R65_)≤ 0.675, and 0.457 ≤HDI(θ_F15R95_)≤ 0.773, fig. S1, C to I). Together, these observations indicate a strong transgenerational response of *P. pacificus* RSC011 to the *Novosphingobium* diet that results in multigenerational expression of the predatory mouth form.

### TEI of the predatory mouth form requires a five-generation exposure

Next, we wanted to know the shortest duration of the exposure to a *Novosphingobium* diet that would induce the TEI of the predatory mouth form. Exposure of RSC011 to *Novosphingobium* for one, two, or three generations did not result in TEI of the Eu phenotype (Fig.2, A to D). When we exposed naïve RSC011 worms to *Novosphingobium* for four generations, we observed intergenerational inheritance that lasted for two generations (Fig. 2E). However, the mean probability of developing the Eu mouth form in the F4R3 generation dropped below 0.5 (θ̄_F4R3_ =0.385, 0.21 ≤HDI(θ_F4R3_)≤ 0.565) (Fig.2E). In contrast, a five-generation exposure resulted in a mean probability of developing the Eu mouth form of 0.514≤θ̄ ≤0.83 for four generations when reverted back to *E. coli* (Fig.2F). Thus, TEI of the predatory mouth form builds up after multiple generations and requires an exposure of at least five generations on *Novosphingobium*. These findings are different from most of the studies in *C. elegans* that observe TEI already after transient environmental exposure (*12*, *16*, *53*). Therefore, our results describe a new type of TEI that requires multigenerational induction. Note that such dietary effects can only be observed in long-term environmental induction experiments.

**Fig. 2.**
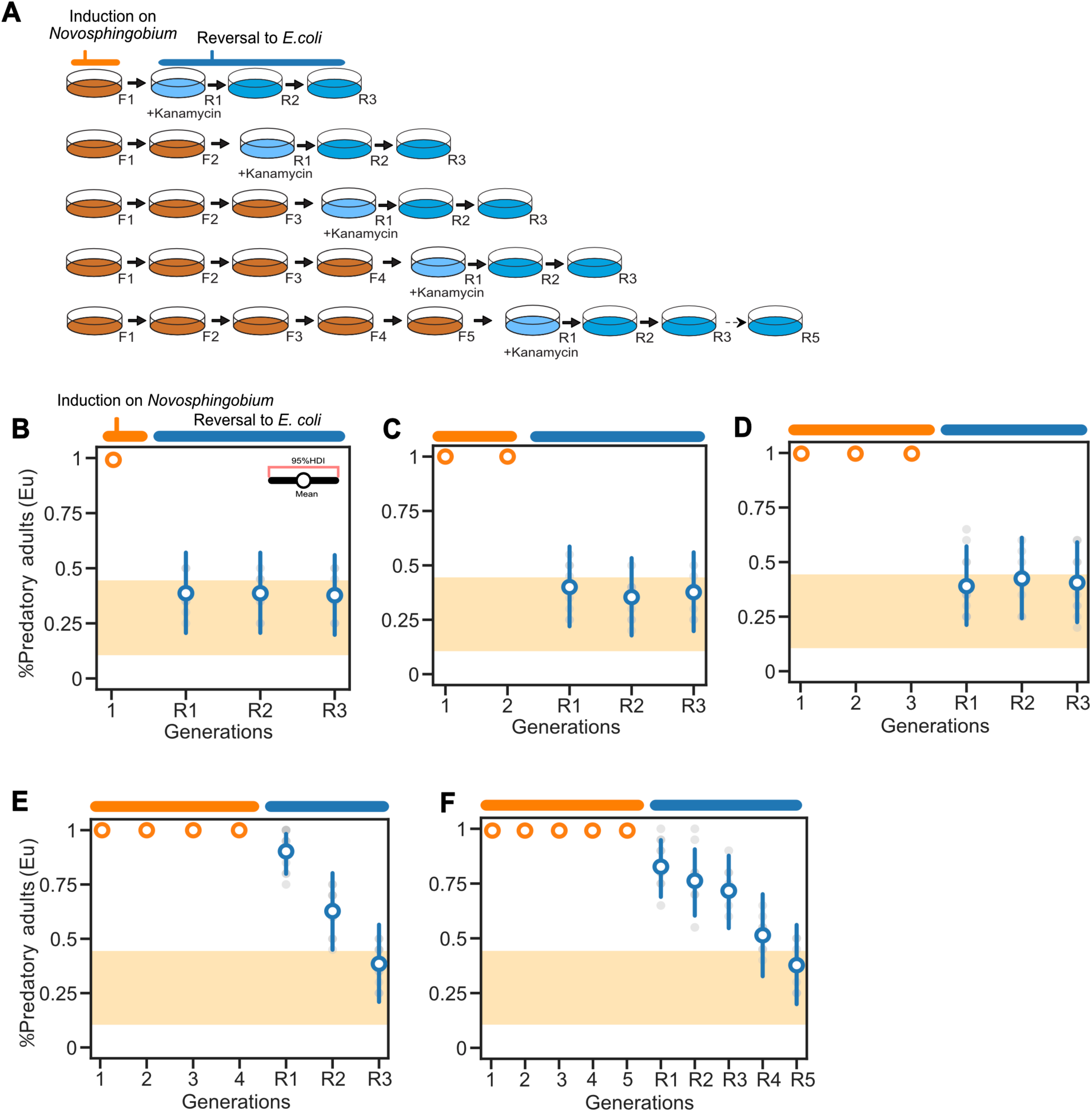
Transgenerational inheritance of the predatory mouth form requires a minimum of five-generation exposure. (**A**) Graphical depiction of the tracking of the predatory morph responses after exposure to *Novosphingobium* (orange) for a certain number of generations and subsequent reversal to *E. coli* (blue). (**B**) Mean probability of predatory mouth-form after exposure to *Novosphingobium* for one generation, (**C**) two generations, (**D**) three generations, (**E**) four generations, and (**F**) five generations following reversal to *E. coli*. 95% HDI estimates of the Eu mean that overlap with the upper limit of the baseline response (yellow) are considered not to be credibly different from the control on *E. coli*. Orange and blue ovals above B to F highlight generations on *Novosphingobium* and *E. coli*, respectively. Yellow region shows the RSC011 baseline response averaged across 101 generations on *E. coli*.

### Forward genetic screening identifies mutants defective in memory transmission of the predatory mouth form

To unravel the molecular machinery involved in the TEI of the predatory mouth form, we coupled the long-term environmental induction experiment with unbiased forward genetic screening for TEI-defective mutants. Specifically, we mutagenized 100 hermaphrodites that were grown on *Novosphingobium* for 14 generations with ethylmethylensulfate (EMS) (Figure 3A).

**Fig. 3.**
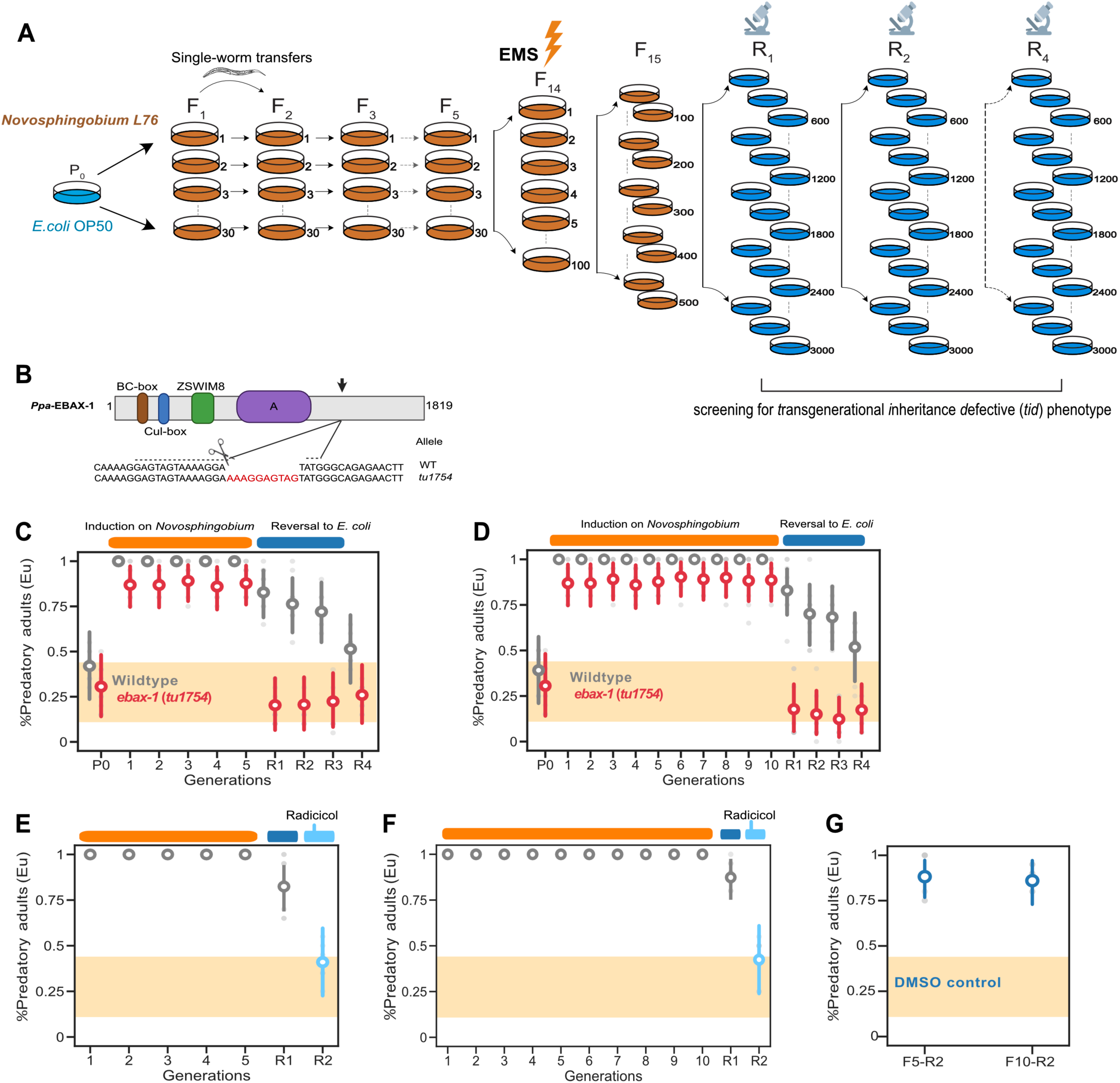
Long-term environmental induction combined with forward genetic screening identifies regulators of transgenerational inheritance. **(A**) Graphical representation of the EMS mutagenesis experiment performed on the F14 generation on a *Novosphingobium diet*. Screening for *transgenerational inheritance defective* (*tid*) mutants was performed after reversal to *E. coli*, and mutant candidates were re-screened in subsequent generations. Some icons are from Biorender.com. (**B**) Domain composition of the substrate-receptor E3 ubiquitin ligase *Ppa-*EBAX-1 that consists of 1819 amino acids. We generated CRISPR/Cas9-induced mutation in the predicted intrinsically disordered region (IDR) of *Ppa-*EBAX*-1* as highlighted (arrow). Insertion of a 10 bp fragment results in a frameshift allele. (**C**) Mean probability of the predatory mouth form in *Ppa-ebax-1* mutant animals upon *Novosphingobium* exposure for 5 generations, and (**D**) 10 generations and reversal to *E. coli*. (**E**) Mean probability of the predatory morph upon radicicol treatment during reversal (R2) to *E. coli* after 5 generations and (**F**) 10 generations of *Novosphingobium* exposure. (**G**) Mean probability of the predatory morph on *E.coli* supplemented with DMSO as controls for the radicicol supplementation experiment. Orange and blue ovals above C to F indicate generations on *Novosphingobium* and *E. coli*, respectively. Light blue ovals above E to F show generations exposed on E. coli supplemented with radicicol. The 95% HDI for the means are based on Bayesian model and shows the probability of developing the Eu morph inferred from the experimental data. Yellow region highlights the RSC011 baseline response averaged across 101 generations on *E. coli*.

After the F15 generation on *Novosphingobium*, we singled out 3,000 F15R1 animals that harbor potential homozygous mutations at individual loci, on kanamycin plates with kanamycin-resistant *E. coli*. We screened for mutants that would no longer show TEI of the predatory mouth form and instead, have low probability of developing the Eu mouth form (θ) on *E. coli* similar to naïve worms (0.115 ≤HDI(θ_Control_)≤ 0.436) (Fig.3A). After screening the F15R1 and re-screening the F15R2 to F15R4 generations, a total of 75 mutants were found with a *transgenerational inheritance defective* (*tid*) phenotype and a 20-40% Eu mouth-form ratio. We performed whole genome sequencing of these mutants and searched for genes with multiple independent alleles. We set an arbitrary threshold of three alleles to consider a gene as a candidate based on previous experience (*46*, *54*). Indeed, we found mutations in *Ppa-ebax-1* and *Ppa-daf-21*/Hsp90 in three independent *tid* alleles (table S1). Both genes are 1:1 orthologous to the corresponding genes in *C. elegans* (fig. S3, A to B), where they have been implicated in several biological processes (*55–59*).

### Substrate-receptor E3 ubiquitin ligase EBAX-1 is required for the TEI of the predatory morph

The identification of the *E*longin *B*C-binding *ax*on regulator *ebax-1* as a potential candidate gene involved in TEI of the predatory mouth form was intriguing. In *C. elegans*, *ebax-1* was first identified as part of the protein quality control mechanism that guards the SAX-3/Robo receptor in developing neurons (*58*). While this original study had suggested an interaction of EBAX-1 with the heat shock protein DAF-21/HSP90 in *C. elegans*, more recent studies indicated that *Cel-*EBAX-1, like its human and *Drosophila* homologs ZSWIM8 and *Dora*, is also involved in microRNA (miRNA) turnover (*55–57*). These studies suggested a novel type of miRNA regulation, referred to as ‘target-directed miRNA degradation’ (TDMD). Current models assume that after the pairing of an mRNA or lncRNA trigger with an Argonaute (Ago)-miRNA complex, a ubiquitin ligase will be recruited, causing the ubiquitination and proteasomal degradation of the Ago protein (Fig. 4A) (*55*, *56*, *60*). Subsequently, the miRNA is degraded by cellular nucleases, resulting in a reversed regulatory outcome from canonical miRNA-mediated gene silencing (*61*, *62*). In mice, ZSWIM8 is known to destabilize many miRNAs that occur in miRNA clusters (*60*). In addition to EBAX-1/ZSWIM8, also DAF-21/Hsp90 has been reported to be involved in miRNA regulation, i.e. the possible loading of highly abundant miRNAs of the *miR-35* family into Ago proteins in *C. elegans* oocyte (*63*, *64*). The isolation of *Ppa-ebax-1* mutants in our screen for a *tid* phenotype could suggest a role of ubiquitin ligase and miRNA signaling in the transmission of epigenetic memory in *P. pacificus*.

**Fig. 4.**
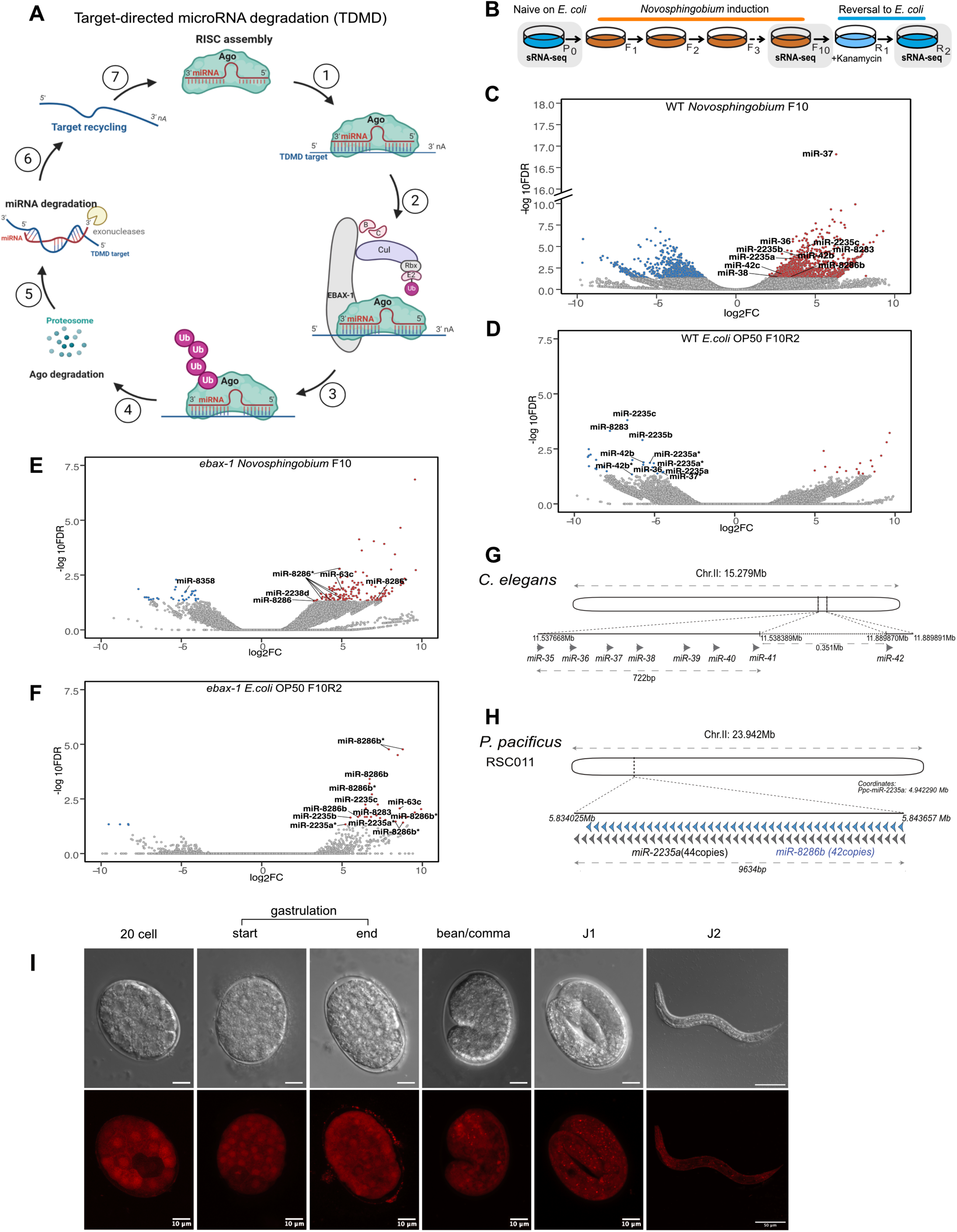
*Ppa-ebax-1* destabilizes levels of *miR-2235*/*miR-35* family in *P. pacificus.* (**A**) Molecular model for target-directed miRNA degradation (TDMD) based on recent studies in mammals and flies. Extensive miRNA-target pairing is recognized by the ZSWIM8/EBAX-1 ubiquitin ligase, and results in subsequent polyubiquitination of the Argonaute (AGO) protein, leading to Ago proteolysis and miRNA decay. Created with Biorender.com. (**B**) Experimental design for small RNA sequencing before, during and after *Novosphingobium* induction. Worm generations used for sequencing are highlighted in gray. (**C**) Small RNA sequencing of lines exposed to *Novosphingobium* for 10 generations compared with naive unexposed animals (n=2 biological replicates for F10 *Novosphingobium*, 3 biological replicates for P0). (**D**) Small RNA sequencing of worms after reversal to *E. coli* for two generations compared with lines exposed to *Novosphingobium* for 10 generations (n=2 biological replicates for each condition). (**E**) Small RNA sequencing of *Ppa-ebax-1* mutants compared to wild type controls, both exposed to *Novosphingobium* for 10 generations (n=2 biological replicates for each condition). Asterisk *, miRNA isoforms. (**F**) Small RNA sequencing of *Ppa-ebax-1* mutant lines compared to wild type controls after reversal to *E. coli* for two generations (n=2 biological replicates for each condition). Asterisk *, miRNA isoforms. (**G**) A schematic representation of the *miR-35* family members in *C. elegans* (*miR-35* to *miR-41*). Grey arrow, *miR-35* members. (**H**) A schematic representation of the heavily expanded *miR-2235a/miR-35* locus *in P. pacificus*. Grey arrow, *miR-2235a*, blue arrow, *miR-8286b*. (**I**) The *miR-2235a* promoter region drives ubiquitous expression during embryogenesis and early juvenile stages. *Ppa-miR-2235a*p::TurboRFP transcription reporter construct is observed starting from 20 cell stage throughout the embryo.

To start testing these hypotheses, we first generated CRISPR alleles of *Ppa-ebax-1* in the RSC011 background, one of which has a frameshift mutation (*Ppa-ebax-1(tu1754)*) (Fig. 3B). While this allele results in a premature stop codon, two other alleles, *Ppa-ebax-1(tu1755)* and *Ppa-ebax-1(tu1756),* are in-frame deletions of 60bp and 12 bp, respectively. All three alleles are viable and grow normal on *E. coli*; however, the *Ppa-ebax-1(tu1754)* allele resulting in a frameshift has locomotion defects and a strong egg-laying-defective phenotype similar to *Cel-ebax-1* mutants (fig. S3, F to H). When grown on an *E. coli* diet, all three *Ppa-ebax-1* mutants have an Eu mouth-form ratio similar to wild type animals (Fig. 3C and fig. S3, I and J).

Similarly, all three *Ppa-ebax-1* alleles are strongly but not completely induced on the *Novosphingobium* diet with the mean probability of developing the Eu mouth form 0.859≤θ̅*_tu_*_1754_≤0.916 (Fig. 3C and fig. S3, I and J). In contrast, all three mutant lines returned immediately to the baseline Eu mouth-form ratio of RSC011 (0.113 ≤HDI(θ_R1(tu1754)_)≤ 0.423) after food reversal (Fig. 3, C and D and fig. S3, I and J). Thus, these experiments indicate a role for *Ppa-*EBAX-1 in transgenerational inheritance of the predatory mouth form and provide the first evidence of a role of ubiquitin ligases in memory transmission.

### The *daf-21*/Hsp90 inhibitor radicicol eliminates TEI of the predatory mouth form

Next, we tested if *Ppa-daf-21/hsp90* is also involved in TEI of the Eu mouth form. As previous work in the *P. pacificus* wild type strain PS312 had indicated that CRISPR-induced frameshift mutations in *Ppa-daf-21/hsp90* resulting in *loss-of-function* alleles are lethal (*59*), we tried to rebuild the amino acid substitutions in *Ppa-daf-21* obtained after EMS mutagenesis using CRISPR editing. However, we were unsuccessful to rebuild such mutations and all obtained frameshift mutations in *Ppa-daf-21* were lethal. Therefore, we used the HSP90 inhibitor ‘radicicol’ that is known to specifically target only HSP90 family members in many organisms including *P. pacificus* (*59*). We grew wild type RSC011 animals on *Novosphingobium* and added the radicicol inhibitor during the reversal to *E. coli* starting in the F5R2 and F10R2 generations, respectively (Figures 3, E and F). In contrast to untreated animals (Fig. 3G), addition of radicicol in the F5R2 generation caused an immediate reduction in the probability of developing the Eu mouth form from θ_R1_ = 0.874 (0.752 ≤ *HDI*(θ_R1_) ≤ 0.975) to θ_R2_ = 0.424 (0.239 ≤*HDI*(θ_R1_) ≤ 0.609) (Fig. 3E). We found a similar reduction in Eu frequency when applying radicicol after an exposure to *Novosphingobium* for 10 generations (Fig. 3F). These results indicate that, similar to *Ppa-*EBAX-1, *Ppa*-DAF-21/HSP90 is required for TEI of the predatory mouth form.

### A *Novosphingobium* diet results in upregulation of the heavily expanded *miR-2235a*/*miR-35* family

Given the role of *Cel-ebax-1* and its *Drosophila* and human homologs in TDMD (Fig.4A) (*55–57*), we wanted to know if *Ppa*-EBAX-1 also affects miRNA expression. Therefore, we first performed small RNA (sRNA) sequencing of wild type animals at three different time points throughout the dietary induction experiment (Fig. 4B). Specifically, we sequenced naïve animals at the beginning of the experiment, the F10 generation on *Novosphingobium* before reversal and the F10R2 generation after reversal back to *E. coli*. To identify diet-sensitive sRNAs, we considered padj< 0.05 for capturing differential expression across conditions. Naïve worms exposed to *Novosphingobium* for 10 generations resulted in the differential expression of 2,644 sRNAs, of which 1,538 were upregulated (Fig. 4C; Data S1A). For the identification of miRNAs in this dataset, we used the previously established catalogue of *P. pacificus* miRNAs that are publicly available at miRBase (*65–70*) (for details see Materials & Methods). Among the up-regulated sRNAs were 30 miRNA species, nine of which belong to the *miR-35* family that is known to be regulated by EBAX-1 in *C. elegans* (Fig. 4C; Data S1A) (*55*, *71*). After the reversal of wild type worms to the *E. coli* diet, there are only 45 differentially expressed sRNAs relative to the *Novosphingobium* F10 generation, the majority of which (26/45) is downregulated (Fig. 4D; Data S1B). This group of downregulated sRNAs contains six of the nine *mir-35* family members that were previously significantly upregulated. Thus, members of the *P. pacificus miR-35* family strongly respond to a change in diet.

In *C. elegans*, the *miR-35* family consists of eight miRNAs (*miR-35 – miR-42*) whose genes are located on a single chromosome with seven of them being clustered within a 700 bp region in an intron of another gene (Fig. 4G) (*71*, *72*). These miRNAs are loaded into the *C. elegans* oocyte where they represent the most abundantly expressed miRNA; however, during embryogenesis these miRNAs are largely degraded (*71*). Deletion of the *miR-35 -miR-41* family members results in strong phenotypic defects, whereas the additional deletion of *miR-42* causes embryonic lethality (*72*, *73*). In *P. pacificus,* several but not all members of the *miR-35* family have been annotated in miRBase. When we mapped the entirety of the *miR-35* family onto the *P. pacificus* genome we found three major differences compared to *C. elegans*. First, *P. pacificus* contains many more members of this miRNA family, several of which have been designated as *miR-2235a*, *miR-2235b* and *miR-2235c* (fig. S5A). Second, single copies of *miR-35* family members are found on all but one chromosome (fig. S5A). Most strikingly however, one locus in the *P. pacificus* RSC011 genome contains 44 identical copies of *mir-2235a* on chromosome II (Fig. 4H). These copies are evenly spaced with a nearly identical spacer of 202 bp. Intriguingly, we observed another miRNA, *miR-8286b*, from a different miRNA family in this spacer with a total of 42 identical copies (Fig. 4H). As a result, this locus of approximately 10 kb in length contains more than 80 miRNAs in an otherwise gene desert (Fig. 4H). Note that in contrast to *miR-2235a,* there are no other copies of *miR-8286b* elsewhere in the *P. pacificus* RSC011 genome. Interestingly, levels of *miR-8286b* are also significantly upregulated when worms are grown on *Novosphingobium* (Fig. 4C; Data S1A). Genome-wide analysis revealed that the *miR-2235a* cluster on chromosome II is the largest cluster detected in the *P. pacificus* genome and is unique in harboring identical copies of the same miRNA and containing another interspersed miRNA in the spacer region (fig. S5B). Note that a recent developmental small RNA transcriptomic study in *P. pacificus* found that *miR-2235a* is one of the most abundantly expressed miRNAs and similar to *C. elegans,* its expression ceases during early development (*74*). Taken together, dietary changes in *P. pacificus* result in differential expression of sRNAs including a heavily expanded family of miRNAs that in *C. elegans* is destabilized by the ubiquitin ligase EBAX-1 (*55*).

### Absence of *Ppa-ebax-1* results in the upregulation of the *miR-2235a* family

Next, we carried out sRNA sequencing of *Ppa-ebax-1(tu1754)* mutant animals at the same three time points during dietary induction and compared differential expression of sRNAs to that of wild type animals. Growing *Ppa-ebax-1(tu1754)* mutants on *Novosphingobium* resulted in 206 sRNA being differentially expressed relative to wild type animals (Fig. 4E; Data S1C). Of these 206 sRNAs, 172 were significantly upregulated in *Ppa-ebax-1(tu1754)* mutants. All *miR-35* family members, including *miR-2235a*, that were significantly upregulated on *Novosphingobium* in the wild type background, are similarly upregulated in *Ppa-ebax-1(tu1754)* mutant animals (Fig. 4E; Data S1C). In contrast, after reversal to *E. coli*, in the F10R2 generation, there are 35 sRNAs that are differentially expressed in *Ppa-ebax-1(tu1754)* mutants relative to wild type (Fig. 4F; Data S1D). Of these 35 sRNAs, 31 are significantly upregulated, including three members of the *miR-35* family with the exact mature sequence of *miR-2235c-3p*, *miR-2235b-3p*, *miR-8283-3p* (Fig. 4F; Data S1D).

Recent studies in mice and flies indicated that miRNA processing can result in trimmed and tailed miRNA versions (*75*). To capture tailing and trimming events at the 3’end of miRNAs during *Novosphingobium* exposure and reversal to *E. coli*, we used the first 18 nt of the mature miRNA sequence in the *miR-2235a* locus and combined all miRNA isoforms into single counts. Indeed, we found 10 miRNA variants associated with the *miR-2235a* locus are also significantly upregulated in *Ppa-ebax-1(tu1754)* mutants relative to wild type (Fig. 4F; Data S1D). This includes two variants that are longer (<23 nt) and shorter (21 nt) versions of *miR-2235a*, and eight variants of *miR-8286b* that are longer 3’ isoforms (20-26 nt long) (Fig. 4F; Data S1D).

These variants contain nucleotides that do not exist in the parental gene sequence and can therefore be considered as non-templated (*76*). Further analysis revealed that the tailed and trimmed versions of the mature *miR-2235a* and *miR-8286b* are upregulated upon the loss of *Ppa-ebax-1* (fig. S4, A and B). Finally, the 22-nt mature *miR-2235a* and 19-nt mature *miR-8286b* miRNAs are also upregulated, but not significantly (padj>0.05). Together, these findings indicate that the loss of *Ppa-ebax-1* results in the upregulation of several *miR-2235/miR-35* family members and destabilizes miRNAs associated with the highly expanded and repetitive *miR-2235a* locus. In addition, the observation of non-templated nucleotide additions and removals at the 3’-end of *miR-2235a* and *miR-8286b* might suggest that they are subject to TDMD or TDMD-like processes. This would be consistent with a repressive role of the *miR-2235a* cluster in TEI of the predatory mouth form.

The identification of primary triggers of EBAX-1/ZSWIM8/Dora-mediated miRNA degradation has been shown to be challenging (*77*). Work in vertebrates so far identified only four(*78–80*), whereas studies in *Drosophila* found six endogenous RNA triggers (*57*). While TDMD-inducing targets usually exhibit extended base pairing with the miRNA 3’ end (*55–57*), studies in *C. elegans* suggested that for the decay of *miR-35* family members the seed sequence of the miRNA is sufficient (*71*). Therefore, it is possible that mechanisms involved in memory transmission of the predatory mouth form in *P. pacificus* might not require extensive pairing beyond the 3’-end. We used existing bioinformatic pipelines to search for potential triggers (*77*) and found hundreds of predicted matches with full or partial complementarity to *miR-2235a* in the *P. pacificus* genome (fig. S5C; Data S3). Not surprisingly, relaxation of sequence complementary from 8mer to 6mer increased the number seed matches in predicted 3’ UTR binding sites (fig. S5C). However, we did not identify matches with known genes in the mouth-form GRN. Note that a similar analysis with *miR-8286b* showed higher numbers of potential 3’ UTR binding sites; while the number of potential matches with extended complementarity (TDMD) is lower for *miR-8286b* (fig. S5C; Data S3).

To study the expression of the miRNAs in the *miR-2235a* cluster, we generated a reporter construct using a 1.4 kb fragment upstream of the first copy of *miR-2235a* in the cluster fused to a *Pristionchus*-codon optimized TurboRFP (*81*). We detected strong zygotic expression in early embryogenesis similar to *miR-35* expression in *C. elegans* (*82*)(Fig. 4I.). In contrast to *C. elegans*, the RFP expression persisted throughout the J1 stage, which in *P. pacificus* remains in the egg, and the J2 stage after hatching (Figure 4I). However, there is no detectable expression in later post-embryonic stages, consistent with the miRNA abundance reported in developmental small RNA transcriptomics (*74*). Similarly, a *Ppa-ebax-1* transcriptional promoter construct showed a dynamic embryonic and juvenile expression starting from the comma-stage and peaking in the J1 stage with expression in the pharynx and tail region (fig. S3C). In adult stages, *Ppa-ebax-1*p::TurboRFP is expressed in pharyngeal neurons, the vulval region and in the gonadal tissues (fig. S3, D and E).

### Deletions of the *miR-2235a* locus are viable

Given the upregulation of *miR-2235a* in *Ppa-ebax-1* mutants and the tailing and trimming of miRNAs of the *miR-2235a* cluster, we speculate that *miR-2235a* may play a repressive role in the TEI of the predatory mouth form. However, to the best of our knowledge, miRNAs have so far not been shown to be involved in transgenerational inheritance, which is different from siRNA and piRNAs (*19*). To study if miRNA signaling including *miR-2235a* is involved in the TEI of the predatory mouth form, we targeted the *miR-2235a* locus. Note that targeting the *miR-2235a* locus for deletion by CRISPR is challenging because large-scale deletions of more than 1 kb have not been regularly reported in *P. pacificus* (*81*). Nonetheless, the highly repetitive nature of the 10 kb locus might allow the use of sgRNAs that can bind repeatedly to target sequences in this region. Indeed, we were able to generate a sgRNA that potentially binds to the *miR-2235a* sequence (Fig. 5A). To identify deletions of the majority or even the entire locus, we designed PCR primers in the flanking regions with unique sequence specificity (Fig. 5A). Note that PCR experiments with these primers will not result in PCR products in wild type animals, because standard PCR conditions do not result in the amplification of 10 kb fragments (fig. S5D). In contrast, large deletions in the *miR-2235a* locus might result in smaller fragments that can be amplified. Using this strategy, we isolated two mutant lines out of 200 tested progenies of CRISPR-injected animals with large deletions in the *miR-2235a* locus. Sequencing analysis showed that both mutant lines exhibited nearly a complete deletion of the *miR-2235a* locus (Fig. 5B). Specifically, original sequencing of PCR-amplified products revealed that *miR-2235(tuDf9)* and *miR-2235(tuDf10)* carry deletions of 9,408 and 9,632 bps, respectively, with two and one copies of *miR-2235a* remaining (Fig. 5B). Whole genome resequencing of both mutant lines confirmed the exact size of these deletions and the fact that only one and two copies of the microRNA are still present (Fig. 5C).

**Fig. 5.**
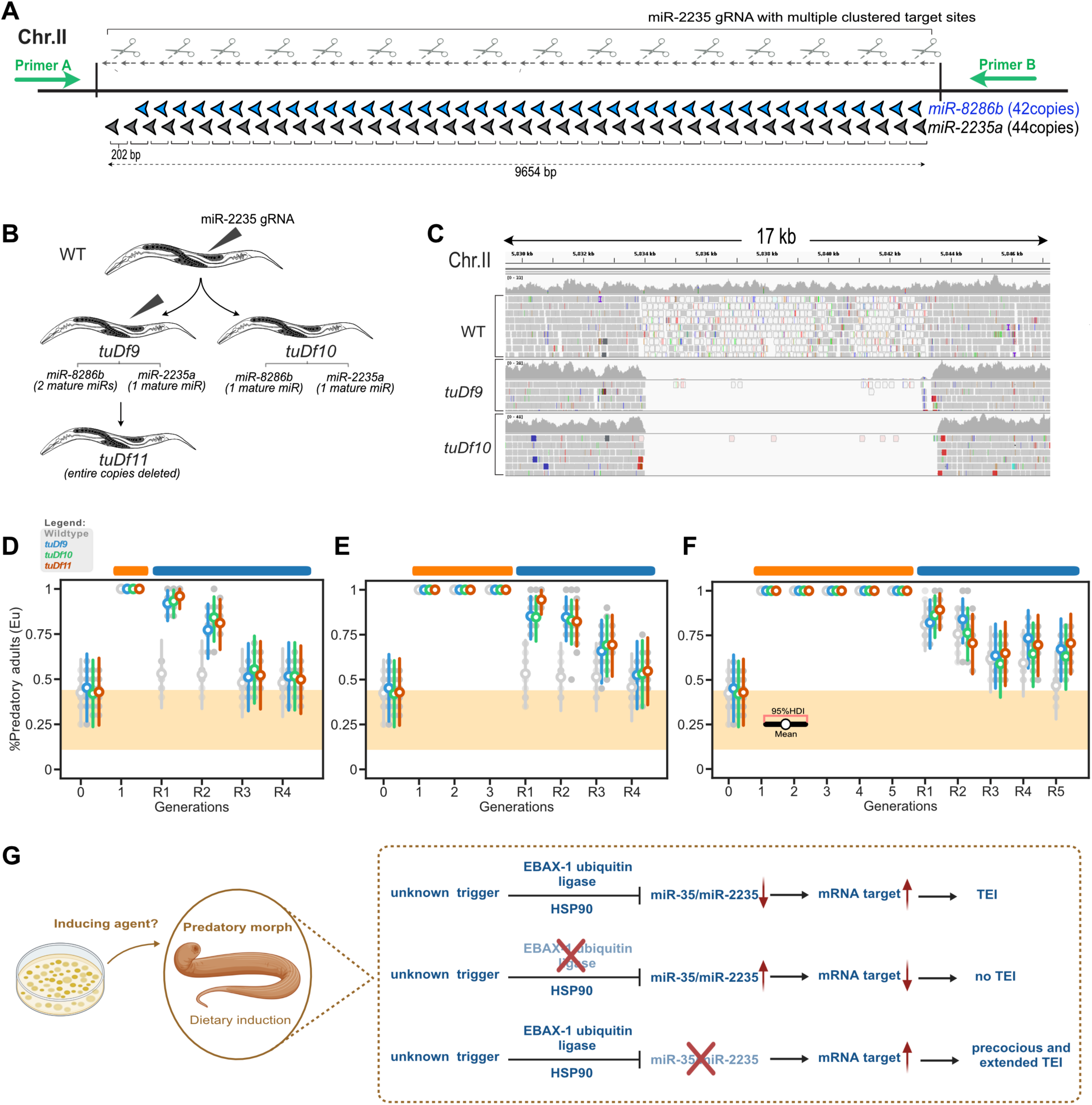
Deletion of the *miR-2235a* locus results in precocious and extended transgenerational inheritance of the predatory mouth form. (**A**) Design of the CRISPR/Cas9 experiment to induce deletions of the *miR-2235a* locus. A sgRNA with multiple binding sites in *miR-2235a* was used. Primers flanking the target locus (green arrows) were designed to amplify shorter fragments as a consequence of large deletions. (**B**) A schematic for the CRISPR/Cas9 induced deletion for all three deficiency alleles. (**C**) Whole genome resequencing confirms large deletions of the *miR-2235a* locus in the homozygous viable mutant lines *tuDf9* and *tuDf10*. (**D**) Mean probability of the predatory mouth form of all deficiency alleles after exposure to *Novosphingobium* for one generation, (**E**) three generations, and (**F**) five generations followed by reversal to *E. coli*. Wiltype, grey; blue, *tuDf9*; green, *tuDf10*; red, *tuDf11.* Orange and blue ovals above D to F indicate generations on *Novosphingobium* and *E. coli*, respectively. The 95% HDI for the means are based on Bayesian model and shows the probability of developing the Eu morph inferred from the experimental data. The yellow region displays the upper and lower limit of the RSC011 baseline averaged across 101 generations on *E. coli*. (**G**) Proposed model depicting transgenerational inheritance of the predatory morph in wild type animals, *ebax-1* and *miR-2235a* alleles as a result of *Novosphingobium* induction. A currently unknown trigger after *Novosphingobium* exposure induces *Ppa-ebax-1* activity leading to *miR-2235a* destabilization. As a consequence of miRNA degradation, target gene/s will be expressed resulting in TEI of the Eu morph.

Both of these deficiencies did not result in the deletion of the entire cluster, maybe because such lines would be embryonic lethal similar to the knockout of *C. elegans miR-35 – miR-42*. To directly test this hypothesis, we performed another CRISPR experiments in the *miR-2235a(tuDf9)* mutant background to delete the remaining miRNA copies using the original sgRNA. Indeed, we were able to isolate a mutant, *miR-2235(tuDf11)*, in which the entire *miR-2235a* cluster is deleted (Fig. 5B). *miR-*2235 (*tuDf11*) mutant animals are viable and phenotypically wild type with no obvious growth defects when cultured on a *E. coli* or *Novosphingobium* diet. In the following we used all three deficiencies at the *miR-2235a* locus to study a potential role of *miR-2235a* in transgenerational inheritance.

### Deletions of the *miR-2235a* cluster cause precocious and extended transgenerational inheritance

If *miR-2235a* expression would repress TEI of the Eu mouth form after food reversal, the deletion of the *miR-2235a* locus might result in an earlier onset of TEI and/or an extension of TEI after reversal. Therefore, we used all three *miR-2235a* alleles and reverted them back to *E. coli* after one and three generation exposures to *Novosphingobium*. In addition, we reverted mutants after five generations to investigate potential longer durations of TEI of the predatory mouth form. Indeed, we found that all three *miR-2235a* mutants show precocious and extended transgenerational inheritance. Specifically, in the F1R1 and F1R2 generations, mutants exhibit an elevated probability of developing the predatory mouth form (0.892 ≤ HDI5θ_F1R1-*tuDf11*_6 ≤ 1.0, and 0.664 ≤ HDI 5θ_F1R2-*tuDf11*_6≤ 0.94). However, the F1R3 generation (0.334 ≤HDI 5θ_F1R3-*tuDf11*_6 ≤ 0.711) was not credibly different from the base line (0.115 ≤ HDI(θ_Control_) ≤ 0.436) (Fig. 5D). Thus, a one-generation exposure to *Novosphingobium* results in intergenerational inheritance of the Eu morph that lasts for two generations. In contrast, reversal after three generation exposure to *Novosphingobium*, results in precocious transgenerational inheritance with (0.862 ≤ HDI(θ_F3R1-*tuDf11*_) ≤ 0.999, 0.687 ≤ HDI(θ_F3R2_) ≤ 0.941, and 0.517 ≤ HDI(θ_F3R3_) ≤ 0.858) (Fig. 5E). In contrast, wild type animals showed no memory after such early reversals as shown previously (Fig. 5, D and E). When we reverted *miR-2235a* mutants back to E*. coli* after 5 generations, we observed TEI of the Eu mouth form lasting for 5 generations (0.533 ≤ HDI(θ_F5R5-*tuDf11*_) ≤ 0.869) (Fig. 5F), whereas wild type animals returned to baseline level lower than 50% Eu in the F5R5 generation (Fig. 5F). Taken together, these experiments support a role of *miR-2235a* as a repressor of epigenetic memory with the increased removal of *miR-2235a* copies resulting in precocious inter- and transgenerational inheritance.

## Discussion

This study establishes a synthesis of multiple novel approaches to investigate transgenerational inheritance. Conceptually, the coupling of long-term environmental induction experiments with the analysis of a natural, phenotypically plastic trait and the subsequent involvement of forward genetic analysis revealed mechanistic insight into the machinery involved in memory transmission. This methodology will allow a detailed investigation of associated processes and signifies *P. pacificus* mouth-form plasticity as a new model to investigate memory transmission. The use of forward mutagenesis provides an unbiased approach to identify genes involved in the transmission of non-genetic information across generations. The experiment described in this study used 3,000 ‘F2’ progeny of mutagenized animals and is thus, far from saturation. However, saturation mutagenesis is feasible in *P. pacificus* because, of the short generation time and easy husbandry similar to *C. elegans*. In addition, this study relies on a binary readout that builds on the developmental plasticity of feeding structures. This feeding structure plasticity is not limited to *P. pacificus*, but is a specific characteristic of the majority of nematodes of the Diplogastridae family to which *P. pacificus* belongs (*83*). Indeed, it is the diversity of feeding structures that has resulted in an extraordinary ecological diversification of these nematodes, highlighting the importance of plasticity as facilitator for evolutionary change (*83–85*). Therefore, this study provides important novel findings for developmental plasticity, associated TEI, and the potential role of TEI for evolution.

Mutant analyses described in this work indicate that *Ppa-ebax-1* and the *miR-2235a* locus are involved in TEI of the Eu mouth form after induction through a *Novosphingobium* bacterial diet. One way to explain these findings is that the ubiquitin ligase is required for transgenerational inheritance by destabilizing miRNAs in *P. pacificus*, which would otherwise repress the TEI of the predatory morph. We speculate that this process involves TDMD or a TDMD-like mechanism by targeting Ago protein/s which are core components of the RNA-induced silencing complexes (RISCs) (Fig. 5G). Specifically, a currently unknown trigger, after exposure to *Novosphingobium*, induces *Ppa-*EBAX-1 activity on an Ago-*miR-2235a* complex, which in wild type animals will cause the degradation of Ago protein(s) and the destabilization of *miR-2235a*. This will result in the expression of target gene(s) that will lead to TEI of the Eu mouth form. In *Ppa-ebax-1* mutants, the Ago-*miR-2235a* complex is no longer destroyed, target gene(s) is/are repressed and TEI of the predatory mouth form does not occur (Fig. 5G).

Consistent with this model, the large or complete deletion of the *miR-2235a* locus results in precocious and extended TEI of the predatory morph, whereas mutations in *Ppa-ebax-1* result in the absence of memory. However, it is important to note that the extent to which the observed phenotypes of *Ppa-ebax-1* mutants and deletions of the *miR-2235a* locus are due to processes involving TDMD with extended pairing of the miRNA and its trigger, remains currently unknown. Formally, we cannot rule out alternative models, in which *Ppa-*EBAX-1 and *miR-2235a* are involved in different molecular processes associated with TEI of the Eu mouth form. Indeed, EBAX-1/Dora/ZSWIM8 are known to be involved in mechanisms of miRNA regulation other than TDMD (*57*, *71*).

While *P. pacificus* shares with *C. elegans* the viability of *ebax-1* mutants, it displays one genomic feature that partially limits some aspects of future analysis. *C. elegans* contains two copies of many major Ago genes, i.e*. alg-1/2*, *alg-3/4* and *prg-1/2*. Although the reasons for these gene duplications are unknown, this ‘double gene set’ has resulted in single mutants in these genes being viable in *C. elegans* (for review see (*19*)). In *P. pacificus*, in contrast, only one copy of *alg-1/2*, *alg-3/4* and *prg-1/2* exists, similar to the miRNA-Ago complex cofactor *ain-1/2* known as GW182 in mammals (*86*). Any attempt to knockout *Ppa-alg-1/2* or *Ppa-ain-1/2,* was unsuccessful as homozygous mutants are lethal. This genomic composition limits the analysis of Ago proteins and their involvement in miRNA signaling and transgenerational inheritance in *P. pacificus*.

In conclusion, transgenerational phenomena come in different forms and flavors. Many of them last for three to five generations, whereas others have been reported to last for more than 20 or even 300 generations (*7*, *16*, *87*, *88*). Some cases show stochastic inheritance, others exhibit distinct segregation patterns that have been described to follow major rules (*13*). The molecular machinery involved in TEI points towards an important role of small RNA molecules, Argonaute proteins and also histone modifications. Already early studies suggested a key role of piRNAs (*7*) in *C. elegans*; however, a role of miRNAs have previously not been documented to regulate TEI. Our findings demonstrate a unique example of developmental plasticity in an ecologically-relevant trait that is accompanied by TEI of previous environmental information.

These findings and the subsequent genetic and molecular investigations demonstrate a role of miRNAs in TEI. Together, this work provides a platform and adds a novel model system to better understand the importance of developmental plasticity and transgenerational memory both from a mechanistic perspective and in its evolutionary context.

## Acknowledgments

We would like to thank Heike Hausmann for assisting in freezing nematode cultures. We are also grateful to Drs. Adrian Streit and Catia Igreja for discussions and helpful comments on the manuscript and Dr. Mohannad Dardiry for conceptual discussions at the beginning of this long-term project.

## Author contributions

Conceptualization: SPQ, RJS

Investigation: SPQ

Formal Analysis: SPQ

Methodology: SPQ, AK, CR, RJS

Writing – Original Draft: SPQ, RJS

Writing – Review & Editing: SPQ, A.K, CR, RJS

Visualization: SPQ

Resources: WR, HW, YW

Funding Acquisition: RJS

Supervision: RJS

## Data and materials availability

*P. pacificus* strains and bacterial isolates generated in this work are freely available through the Lead Contact. All the code and data for Bayesian analysis are available https://github.com/shielapearl18/Bayesian-estimates-of-dietary-induced-responses. Sequencing data that was generated for the current study has been submitted to the European Nucleotide Archive under the project accession PRJEB74486. All data are available in the main text or the supplementary materials.

## Supplementary Materials

Figs. S1 to S5

Tables S1 to S2

References (89–119)

Data S1 to S3

## Materials and Methods

### Nematode culture conditions

The ancestral *P. pacificus* RSC011 isolate used in this study was frozen within the first 5-10 generations after original isolation from Coteau Kerveguen on La Réunion to minimize domestication and thereby, facilitate the investigation of diet-induced plasticity and transgenerational epigenetic inheritance. Nematodes were grown under standard nematode growth conditions on NGM plates seeded with either *Escherichia coli* OP50 or *Novosphingobium* L76 and maintained at 20°C. We refrained from exposing the RSC011 stock to all other types of stressors such as bleaching, starvation, extreme temperature fluctuations as they are known to potentially influence mouth-form ratios.

### Bacterial strains conditions

All bacterial strains were grown overnight in LB (Lysogeny broth) supplemented with 50 μg/ml kanamycin where required. Bacteria were grown at 30 °C or 37 °C depending on the species and 6 cm nematode growth medium (NGM) plates were seeded with 300 μl bacterial overnight cultures and were incubated for 2 days.

### Long-term experimental evolution (LTEI) approach

LTEI was performed using the ancestral *P. pacificus* RSC011 strain that is preferentially St (20-40% Eu) on a standard *E. coli* OP50 diet. Using the complete brood of a single hermaphrodite, 110 clonal lines were established by picking single J4 larvae on a *Novosphingobium* L76 diet. J4 worms were initially left for at least an hour on a *Novosphingobium* diet to reduce traces of *E. coli* OP50 before transferring to final F1 *Novosphingobium* plates. All 110 lines were transferred every 4 days to new plates by picking a single J4 animal to start the new generation. Rigorous worm maintenance was carried out for an entire year allowing the completion of 101 generations on *Novosphingobium*. Populations on *Novosphingobium* were periodically frozen every 10 generations (up to F100) and stored in liquid nitrogen which allows accurate evaluation of the evolved states.

### Dietary reversal experiments

Overnight cultures of *E. coli* OP50 and *Novosphingobium* L76 were spread to NGM plates and incubated at RT for 2 days. Reversal experiments to an *E. coli* OP50 diet were performed after exposing of RSC011 worms for 15, 25, 35, 45, 55, 65, 75, 85, 95 generations on *Novosphingobium* L76. Worms were initially transferred from *Novosphingobium* L76 to NGM plates supplemented with 50 μg/ml of kanamycin for one generation, then back to *E. coli* OP50-seeded NGM plates for subsequent generations. Exposure to kanamycin for a single generation to eliminate traces of *Novosphingobium* did not affect mouth-form response. Mouth-form phenotyping was examined for the entirety of the reversal experiments (all 15 generations).

### Mouth-form phenotyping

Mouth-from phenotyping was performed using Zeiss Discovery V.20 stereo microscope (X150 magnification) by observing the nematode buccal cavity based on mouth-form identities previously described (*28*). Final mouth-form frequencies are the mean of at least 10-20 replicates (*i.e.* 110 replicates for the LTEI experiment; 10-20 replicates for genetic mutants, and the radicicol supplementation experiments). In each replicate, 20 animals from a single plate were assayed. For the LTEI experiment, phenotyping was done continuously for 25 generations on *Novosphingobium* and periodically every 5 generations (after F25) until 101 generations for all 110 lines. To ensure continuous tracking, a subset of 30 out of the 110 lines were assayed every generation (F26-F29, F31-F34, etc.).

### Statistical analysis

To estimate the probability of developing the Eu mouth form in *P. pacificus* (θ), we constructed a hierarchical Bayesian model (Figures S1A and S1B)

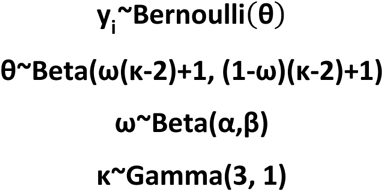

where θ is calculated for each replicate and hyperparameters ω and κ link the biological replicates for a given generation under a given experimental condition (*89*). The model was fitted to the laboratory measurements using PyMC (*90*) in Python 3.11 with Numpy 1.25.2 (*91*). The mean highest density interval (HDI) for θ for a group of observations was used to visualize the inferred probability of developing the Eu mouth form. We ensured that the estimated θ values were stable using common convergence diagnostics with effective sample size (ESS) **≥**10000 (*92*).

### Mutation Rate Calculation

Paired-end Illumina reads were mapped to the RSC011 PacBio genome with bwa (version: 0.7.17-r1188, algorithm: mem)(*93*). Subsequently, the germline short variant discovery workflow (*86*) from the GATK (4.2.5.0) pipeline was employed to mark and remove duplicates (MarkDuplicatesSpark), locally realign reads, call single nucleotide polymorphisms (SNPs) and indels per-sample (HaplotypeCaller) and perform joint calling across multiple samples (GenomicsDBImport and GenotypeGVCFs). SNPs were selected with GATK SelectVariants (--select-type-to-include SNP). Line-specific SNPs were considered to have arisen through mutation accumulation if the alternative allele was homozygous, detected on chromosomes, and met high-quality criteria (DP>=5 and GQ>=20). Mutation rate per line were determined by dividing the number of accumulated mutations in each line by the number of generation (100) and the sample-specific total length of sufficiently covered genomic regions (genome region with DP>=5 in gVCF from GATK HaplotypeCaller for each sample). To mitigate potential biases introduced by problematic genomic regions, number of line-specific SNPs in sliding windows (length: 100bp, step: 20bp) across the chromosomes were calculated and those windows with more than 2 line-specific mutations were excluded from the analysis (*95*).

### EMS mutagenesis

We combined long-term environmental induction to induce heritable mutations in *P*. *pacificus*, and incubated a mixture of *RSC011* J4 larvae grown on *Novosphingobium* L76 for 14 generations in M9 buffer (3 g/L KH_2_PO_4_, 6 g/L Na_2_HPO_4_, 5 g/L NaCl, 1 mM MgSO_4_) with 47 mM ethyl methanesulfonate (EMS) for 4 h (*96*). Subsequently, worms were allowed to recover on agar plates with *Novosphingobium* and 100 actively moving J4 larvae were singled out. After the mutagenized P0 animals had laid approximately 30 eggs, they were killed, and F1 progeny were allowed to develop and reach maturity. A total of 500 ‘F1’ animals were singled out to establish new populations for one more generation on *Novosphingobium* prior to reversal. During reversal to kanamycin plates, we established 3000 ‘F2’ lines (which contained potential homozygous mutants) and allowed to lay ‘F3’ populations for mouth-form screening. Screening was performed for the progenies of ‘F2’ individuals (during reversal) using Discovery V20 stereomicroscope (see above). We searched for lines with a transgenerational inheritance-defective (*tid*) phenotypes, which show an immediate drop of the Eu response to 10-45%, whereas wild-type RSC011 animals keep a mouth-form memory of around 80-100% Eu after reversal. Mutant candidates with a *tid* phenotype and a low Eu mouth-form frequency were transferred to fresh *E. coli* OP50 plates and their progeny were rescreened for additional generations. Out of the 3000 mutagenized F2s progeny, 75 *tid* candidate lines were isolated.

### Whole genome sequencing analysis of mutant lines

Mutant worms were grown on standard NGM plates before DNA extraction. We rinsed plates with M9 buffer and collected worm pellets by slow centrifugation at 1300 rpm for 3 min at 4°C. DNA was extracted with Monarch Genomic DNA Purification Kit following the manufacturer’s protocol. Library preparation and sequencing was done by Novogene. The program BWA mem program (version 0.7.17-r188) (*93*) was used to align raw Illumina reads against the reference genome of the *P. pacificus* RSC011 strain. Initial variant calls were made by the samtools mpileup, bcfutils and varFilter.pl commands (version 0.1.18-r982:295) (*97*). For candidate mutations, we filtered for high-quality, homozygous single nucleotide substitutions that are not shared with resequencing data of the parental strain RSC011. The effect of mutations on protein coding genes was assessed by comparison with the RSC011 gene annotations (*98*). Finally, the number of independent alleles per gene was computed to prioritize candidate genes. Given that classical mapping experiments are unpractical for such an experimental design, all mutant candidate lines were whole-genome sequenced. We considered genes as candidates that had at least 3 independent alleles.

### CRISPR/Cas9-Induced Mutations

Mutations were induced in candidate genes via CRISPR/Cas9 following the protocol described previously (*81*, *99*). Gene-specific sgRNAs and universal trans-activating CRISPR RNA (tracrRNA) were purchased from Integrated DNA Technologies and 5 μl of each 100 μM stock were mixed and denatured at 95 °C for 5 minutes. After allowing the mixture to anneal at RT, Cas9 endonuclease (New England Biolab) was added to the hybridized product and incubated at RT for 5 min. TE buffer was subsequently added to a final concentration of 18.1μM for the sgRNA and 2.5 μM for Cas9. This was injected into the germline of 40-50 *P. pacificus* RSC011 young adults. Eggs from injected P0s were recovered up to 16 h post injection. Post-hatching and 2 days’ growth, these F1 worms were segregated onto individual plates until they had also developed and laid-eggs sufficiently. The genotypes of the F1 animals were further analyzed via Sanger sequencing and mutations were identified before re-isolation of homozygous mutants.

sgRNAs and associated primers utilized in this study can be found in supplementary table. Evidence for the generation of putative null mutants was based upon frame shifts, leading to premature stop codons in the protein coding sequences.

### CRISPR/Cas9-mediated *miR-2235* locus deletion

Large deletions in the *miR-2235a* locus were induced via CRISPR/Cas9 following the protocol indicated above. We used a sgRNA against the *miR-22335a* sequence and injected 60 *P. pacificus* RSC011 young adults. A total of 200 F1 progenies of injected animals were segregated onto individual plates. After egg laying, these F1 animals were used for PCR experiments with primers in the flanking regions of the 10 kb *miR-2235a* locus. Visualizing of PCR product was performed on agarose gel electrophoresis. Individuals with PCR products were considered potential heterozygous candidates for deletions at the locus. We found two potential heterozygous individuals with a ∼ 500 bp PCR fragment. Given that heterozygous and homozygous worms would be indistinguishable based on PCR and Sanger sequencing, we maintained potential heterozygous line by single-worm transfer for several generations and followed sub-lines that gave off clean PCR products in all of the singled progeny. Note that heterozygous animals should segregate 25% wild type animals without a deletion that should result in the absence of the PCR product. The two homozygous mutants were whole genome sequenced (see protocol above), which confirmed original Sanger sequencing results and the deletion at the locus.

### Transgenesis

We generated transcriptional reporter constructs to visualize tissues that actively express *Ppa-ebax-1*. We took 1065 bp upstream sequence of the first Methionine but also extending to the first 515 bp of the gene sequence for optimal primer design. For the *miR-2235a* locus, the 1,455 bp long putative promoter of the first precursor *miR-2235a* sequence was taken. Each of the promoter sequences was PCR-amplified and subsequently assembled into a codon-optimized pUC19 vector containing TurboRFP and the *Ppa-rpl-23* 3’UTR (*81*). Assembly and bacterial transformation was generally done using the NEBuilder HiFi DNA Assembly Kit and NEB-5-alpha *E. coli* cells. Plasmid extraction was done using QIAGEN Plasmid Midi Kit. Linearization of the plasmids was carried out with Pst1 enzyme digestion according to manufacturer’s protocol. Final concentration of the injection mix contained 10 ng/ul of the linearized plasmids with 60 ng/ul Pst1-linearized RSC011 genomic carrier DNA. Flourescent worms were immobilized on a 5% Noble agarose pad with 0.3% NaN_3_ added as an aneasthetic and were screened using a Leica SP8 confocal laser-scanning microscope.

### miRNA target site prediction

The pipeline that searched for TDMD triggers was implemented following previously established in-silico approach in *Drosophila* (*57*). A file annotation for all RSC011 3’ UTRs was loaded as input to start the search for miRNA targets. 3’UTR sequences were derived from RSC011 transcriptome assemblies or from protein homology with the *P. pacificus* PS312 annotations (El Paco V3), by taking the region downstream of the coding sequence within the assembled transcript or 100 bp downstream of the annotated gene. Filtering with the raw RNA-seq data from *ebax-1* animals was implemented, retaining only those reads with more than 10 counts. The input file was queried for binding complementarity towards the 3’region (starting from 13^th^ nucleotide until the last *miR* sequence) of the *miR-2235a* (22 nt-long) and *miR-8286b* (extended to 22 nt-long). Search for complementarity sites to the seed sequence (2-8 nt of the *miR*) was implemented within 30 nt upstream of the 3’ region. Both queries at the seed and 3’end tolerate a maximum of two mismatches.

To search for canonical miRNA-target pairing sites, the query was restricted to the seed sequence of the miRNA. Different types of *miR*-target searches were performed: 8-mer, 7mer, A1-7mer, and 6-mer type of pairing. The search was done independent of any expression dataset. All targets are listed in Data S3.

### Phylogenetic analyses

For *P. pacificus* genes retrieved from the *tid* screen, one-to-one orthologous gene prediction between *P. pacificus* and *C. elegans* were identified from BLASTP searches. One-to-one orthology was further confirmed by phylogenetic analyses. Proteins were then aligned by Clustal Omega (*100*) and output in FASTA format. FASTA files were uploaded to the IQ-TREE tool (*101*, *102*) using default settings with the auto substitution model, and 1,000 bootstrap alignments were calculated under the ultrafast setting. The resulting phylogenetic tree files were visualized using FigTree (http://tree.bio.ed.ac.uk/software/figtree/).

### Radicicol treatment

Radicicol treatment was done as previously described (*59*, *103*). In brief, Radicicol (Sigma-Aldrich) was dissolved in DMSO and then added to NGM agar to the final concentration of 19ug/mL. Solution containing the same amount of DMSO was used as control. Supplementation was done two generations after reversal to *E. coli*.

### RNA sequencing and data analysis

Mixed-staged RSC011 worms grown on *Novosphingobium* and *E. coli* OP50 were collected from three 3 NGM plates. Specifically, worms were pelleted for RNA certain generations of dietary exposure. Total RNA was extracted using Direct-Zol RNA Mini prep kit (Zymo Research) according to the manufacturer’s guidelines. The RNA-seq library preparation and sequencing were done by the company Novogene. Software Hisat2 (*104*) (version 2.1.0) was used to map raw reads to the *P. pacificus RSC011* reference genome, featureCounts (*105*) was used to quantify the counts of reads mapped to the genomic feature based on the RSC011 gene annotations. Tests for differential expression was performed in R (version 4.0.3) using DESeq2 (version 1.18.1) (*106*). Genes with an FDR-corrected *p* value < 0.01 and fold change cutoff of two were considered as significantly differentially expressed. Tests for overrepresentation of protein domains in sets of differentially expressed genes were performed using a Fisher’s exact test in R. We used the FDR method as implemented in the p.adjust function in R to correct for multiple testing and only retained results with adjusted P-value < 0.01.

### sRNA extraction

Total RNA was extracted as described above. After RNA extraction, TAP (*Cap-Clip*™ Acid Pyrophosphatase) was performed by adding a total of 25U TAP to <4 μg total RNA, 5 μl of reaction buffer (10X Cap-Clip™ Acid Pyrophosphatase Reaction Buffer) and PCR water to a total reaction of 50 μl per sample. The reaction was then incubated for two hours at 37°C. RNA clean-up step was done post-incubation using RNA Clean and Concentrator (Zymo Research) as per manufacturer’s recommendations. sRNA-seq library preparation and sequencing were done by the company Novogene with 50-nt single-end reads.

### Analysis of small RNA sequencing data

We trimmed adapters from raw sequencing reads with the software Cutadapt (version 3.5, options: -a AGATCGGAAGA -m 14)(*107*). For each comparison, we counted the number of occurrences for each unique read across the samples (at least two biological replicates per condition). During this step, we discarded reads that were present in only a single sample. DE-seq2 was run to identify significantly differentially expressed sequences between (corrected P-value < 0.05). In order to focus on nematode sequences, we removed differentially expressed sequences that mapped to the genomes of *Novosphingobium* or *E. coli* OP50 by bowtie (version 2.2.3) (*108*). To visualize the distribution of miRNAs across the *P. pacificus* genome, we downloaded 38,589 hairpin sequences from miRBase (release 2023-10-23) which contained 354 sequences from *P. pacificus* (*65–70*, *109*). These sequences were first clustered based on a 90% nucleotide identity cutoff using the cdhit-est program (version 4.8.1, option: -c 0.90) (*110*). We then ran BLASTN (version 2.10.1, option: -evalue 0.001) to search for matches of these sequences in the *P. pacificus* genome (version El Paco) (*111*). BLASTN matches with minimum length of 50 nucleotides and a minimum percentage identity of 90% were then visualized in non-overlapping 100-kb windows.

For combining all miRNA isoforms into single counts, we used string-matching to the first 18 nt of the mature miRNA sequence while also binning reads by lengths from 19–30 nt (*75*).

Trimmed reads were all reads shorter than the mature miRNA length, and tailed reads were all reads longer than the mature length (as indicated in miRBase). In this manner, we were able to visualize the aggregate number of potential trimmed and tailed reads for each miRNA in the heavily expanded *miR-2235a* locus.

### Genome assembly and annotation

Raw PacBio reads of P. pacificus RSC011 were assembled by the software Canu (version 1.4, options: genomeSize=200m -pacbio-corrected) (*112*). For strain RSC011, this resulted in an initial assembly of 54 contigs. This assembly was scaffolded using whole genome sequencing data from recombinant inbred lines of the cross between *P. pacificus* strains RSC011 and RSA076 (*113*). Specifically, whole genome sequencing data were aligned to the raw assembly by BWA aln and sampe programs (version 0.7.17-r188) (*93*). Subsequently, the samtools (version 0.1.18-r982:295) (*97*) mpileup, bcfutils and varFilter.pl commands were run to genotype single nucleotide variant markers between the parental strains RSC011 and RSA076 (*114*). The resulting genotype maps were plotted in R and manually scaffolded after visual inspection. At this step, one contig spanning 4.6 Mb was identified as *E. coli* based on a BLASTN search and was discarded. The resulting genome assembly spans 159.0 Mb of which 156.4 Mb (98.3%) could be placed onto the six chromosomes. To annotate protein-coding genes in the RSC011 assembly, we employed the PPCAC pipeline (version 1) (*115*), which generated evidence-based gene annotations from existing gene annotations for the *P. pacificus* reference strain PS312 (El Paco gene annotations, version 3) (*116*) as well as from RNA-seq data that were generated for this study. We refined the RSC011 gene annotations by removing species-specific orphan genes that overlapped in antisense direction with conserved genes as we previously observed that coding potential on the antisense strand can result in gene annotation errors (*116*, *117*).

Evaluating the completeness of the final RSC011 gene annotations using the BUSCO approach (version 3.1, nematode odb9 data set) (*118*) revealed a completeness level of 89.5%, which demonstrates a comparable degree of completeness with regard to previously assembled *P. pacificus* strain genomes (*117*). Protein domains were annotated by the hmmsearch program with the Pfam-A database of the HMMER package (version 3.3, options: -E 0.001) (*119*)

## References

1. R. C. Painter, C. Osmond, P. Gluckman, M. Hanson, D. I. W. Phillips, T. J. Roseboom, Transgenerational effects of prenatal exposure to the Dutch famine on neonatal adiposity and health in later life. BJOG 115, 1243–1249 (2008).

2. S. Song, W. Wang, P. Hu, Famine, death, and madness: Schizophrenia in early adulthood after prenatal exposure to the Chinese Great Leap Forward Famine. Soc. Sci. Med. 68, 1315–1321 (2009).

3. L. R. Baugh, T. Day, Nongenetic inheritance and multigenerational plasticity in the nematode *C. elegans*. Elife 9, 1–13 (2020).

4. N. Frolows, A. Ashe, Small RNAs and chromatin in the multigenerational epigenetic landscape of *Caenorhabditis elegans*. Philos. Trans. R. Soc. B Biol. Sci. 376, 1–12 (2021).

5. X. Chen, O. Rechavi, Plant and animal small RNA communications between cells and organisms. Nat. Rev. Mol. Cell Biol. 23, 185–203 (2022).

6. F. Santilli, A. Boskovic, Mechanisms of transgenerational epigenetic inheritance: lessons from animal model organisms. Curr. Opin. Genet. Dev. 79, 1–8 (2023).

7. A. Ashe, A. Sapetschnig, E. M. Weick, J. Mitchell, M. P. Bagijn, A. C. Cording, A. L. Doebley, L. D. Goldstein, N. J. Lehrbach, J. Le Pen, G. Pintacuda, A. Sakaguchi, P. Sarkies, S. Ahmed, E. A. Miska, PiRNAs can trigger a multigenerational epigenetic memory in the germline of *C. elegans*. Cell 150, 88–99 (2012).

8. B. A. Buckley, K. B. Burkhart, S. G. Gu, G. Spracklin, A. Kershner, H. Fritz, J. Kimble, A. Fire, S. Kennedy, A nuclear Argonaute promotes multigenerational epigenetic inheritance and germline immortality. Nature 489, 447–451 (2012).

9. S. G. Gu, J. Pak, S. Guang, J. M. Maniar, S. Kennedy, A. Fire, Amplification of siRNA in *Caenorhabditis elegans* generates a transgenerational sequence-targeted histone H3 lysine 9 methylation footprint. Nat. Genet. 44, 157–164 (2012).

10. B. G. Dias, K. J. Ressler, Parental olfactory experience influences behavior and neural structure in subsequent generations. Nat. Neurosci. 17, 89–96 (2014).

11. J. Bozler, B. Z. Kacsoh, G. Bosco, Transgenerational inheritance of ethanol preference is caused by maternal NPF repression. Elife 8, 1–18 (2019).

12. R. S. Moore, R. Kaletsky, C. T. Murphy, Piwi/PRG-1 Argonaute and TGF-β Mediate Transgenerational Learned Pathogenic Avoidance. Cell 177, 1827–1841 (2019).

13. L. Houri-Zeevi, Y. Korem Kohanim, O. Antonova, O. Rechavi, Three Rules Explain Transgenerational Small RNA Inheritance in *C. elegans*. Cell 182, 1186–1197 (2020).

14. M. F. Perez, B. Lehner, Intergenerational and transgenerational epigenetic inheritance in animals. Nat. Cell Biol. 21, 143–151 (2019).

15. O. Rechavi, L. Houri-Ze’evi, S. Anava, W. S. S. Goh, S. Y. Kerk, G. J. Hannon, O. Hobert, Starvation-induced transgenerational inheritance of small RNAs in *C. elegans*. Cell 158, 277–287 (2014).

16. A. Klosin, E. Casas, C. Hidalgo-Carcedo, T. Vavouri, B. Lehner, Transgenerational transmission of environmental information in *C. elegans*. Science 356, 320–323 (2017).

17. N. O. Burton, C. Riccio, A. Dallaire, J. Price, B. Jenkins, A. Koulman, E. A. Miska, Cysteine synthases CYSL-1 and CYSL-2 mediate *C. elegans* heritable adaptation to P. vranovensis infection. Nat. Commun. 11, 1–13 (2020).

18. M. F. Perez, M. Shamalnasab, A. Mata-Cabana, S. Della Valle, M. Olmedo, M. Francesconi, B. Lehner, Neuronal perception of the social environment generates an inherited memory that controls the development and generation time of *C. elegans*. Curr. Biol. 31, 4256–4268 (2021).

19. R. F. Ketting, L. Cochella, Concepts and functions of small RNA pathways in *C. elegans*. Curr. Top. Dev. Biol. 144, 45–89 (2021).

20. L. Houri-Zeevi, G. Teichman, H. Gingold, O. Rechavi, Stress resets ancestral heritable small RNA responses. Elife 10, 1–31 (2021).

21. A. K. Webster, J. M. Jordan, J. D. Hibshman, R. Chitrakar, L. Ryan Baugh, Transgenerational effects of extended dauer diapause on starvation survival and gene expression plasticity in *Caenorhabditis elegans*. Genetics 210, 263–274 (2018).

22. R. Kaletsky, R. Moore, T. Sengupta, R. Seto, C. T. Murphy, Absence of Evidence is Not Evidence of Absence: The many flaws in the case against transgenerational epigenetic inheritance of pathogen avoidance in *C. elegans*. bioRxiv, doi: 10.1101/2024.06.07.597568 [Preprint] (2024).

23. D. P. Gainey, A. V. Shubin, C. P. Hunter, Irreproducibility of transgenerational learned pathogen-aversion response in *C. elegans*. bioRxiv, doi: 10.1101/2024.06.01.596941 [Preprint] (2024).

24. R. E. Lenski, Experimental evolution and the dynamics of adaptation and genome evolution in microbial populations. ISME J. 11, 2181–2194 (2017).

25. R. J. Sommer, L. Carta, S.-Y. Kim, P. W. Sternberg, Morphological, genetic and molecular description of *Pristionchus pacificus* sp. n. (Nematoda: Neodiplogastridae). Fundam. apply. Nematol. 19, 511–521 (1996).

26. N. E. Schroeder, Introduction to *Pristionchus pacificus* anatomy. J. Nematol. 53, 1–9 (2021).

27. G. Bento, A. Ogawa, R. J. Sommer, Co-option of the hormone-signalling module dafachronic acid-DAF-12 in nematode evolution. Nature 466, 494–497 (2010).

28. E. J. Ragsdale, M. R. Müller, C. Rödelsperger, R. J. Sommer, A developmental switch coupled to the evolution of plasticity acts through a sulfatase. Cell 155, 922–933 (2013).

29. J. W. Lightfoot, M. Wilecki, C. Rödelsperger, E. Moreno, V. Susoy, H. Witte, R. J. Sommer, Small peptide-mediated self-recognition prevents cannibalism in predatory nematodes. Science 364, 86–89 (2019).

30. R. J. Sommer, Phenotypic plasticity: From theory and genetics to current and future challenges. Genetics 215, 1–13 (2020).

31. M. Wilecki, J. W. Lightfoot, V. Susoy, R. J. Sommer, Predatory feeding behaviour in *Pristionchus* nematodes is dependent on phenotypic plasticity and induced by serotonin. J. Exp. Biol. 218, 1306–1313 (2015).

32. T. Renahan, R. J. Sommer, Nematode Interactions on Beetle Hosts Indicate a Role of Mouth-Form Plasticity in Resource Competition. Front. Ecol. Evol. 9, 1–9 (2021).

33. M. Ptashne, A genetic switch: Gene control and phage. J. Med. Genet. 24, 789–790 (1986).

34. G. Balázsi, A. Van Oudenaarden, J. J. Collins, Cellular decision making and biological noise: From microbes to mammals. Cell 144, 910–925 (2011)

35. P. M. Brakefield, J. Gatest, D. Keyst, F. Kesbeke, P. J. Wijngaarden, A. Monteiro, V. French, S. B. Carrollt, Development, plasticity and evolution of butterfly eyespot patterns. Nature 384, 236–242 (1996).

36. M. R. Kieninger, N. A. Ivers, C. Rödelsperger, G. V. Markov, R. J. Sommer, E. J. Ragsdale, The Nuclear Hormone Receptor NHR-40 Acts Downstream of the Sulfatase EUD-1 as Part of a Developmental Plasticity Switch in *Pristionchus*. Curr. Biol. 26, 2174–2179 (2016).

37. L. T. Bui, N. A. Ivers, E. J. Ragsdale, A sulfotransferase dosage-dependently regulates mouthpart polyphenism in the nematode *Pristionchus pacificus*. Nat. Commun. 9, 1–10 (2018).

38. B. Sieriebriennikov, N. Prabh, M. Dardiry, H. Witte, W. Röseler, M. R. Kieninger, C. Rödelsperger, R. J. Sommer, A developmental switch generating phenotypic plasticity is part of a conserved multi-gene locus. Cell Rep. 23, 2835–2843 (2018).

39. B. Sieriebriennikov, S. Sun, J. W. Lightfoot, H. Witte, E. Moreno, C. Rödelsperger, R. J. Sommer, Conserved nuclear hormone receptors controlling a novel plastic trait target fast-evolving genes expressed in a single cell. PLoS Genet. 16, 1–27 (2020).

40. S. Sun, H. Witte, R. J. Sommer, Chitin contributes to the formation of a feeding structure in a predatory nematode. Curr. Biol. 33, 15–27 (2023).

41. S. Casasa, E. Katsougia, E. J. Ragsdale, A Mediator subunit imparts robustness to a polyphenism decision. Proc. Natl. Acad. Sci. U. S. A. 120, 1–8 (2023).

42. N. A. Levis, E. J. Ragsdale, A histone demethylase links the loss of plasticity to nongenetic inheritance and morphological change. Nat. Commun. 14, 1–13 (2023).

43. M. S. Werner, T. Loschko, T. King, S. Reich, T. Theska, M. Franz-Wachtel, B. Macek, R. J. Sommer, Histone 4 lysine 5/12 acetylation enables developmental plasticity of *Pristionchus* mouth form. Nat. Commun. 14, 1–14 (2023).

44. T. Theska, R. J. Sommer, Feeding-structure morphogenesis in “rhabditid” and diplogastrid nematodes is not controlled by a conserved genetic module. Evol. Dev. 26, 1–18 (2024).

45. M. S. Werner, B. Sieriebriennikov, T. Loschko, S. Namdeo, M. Lenuzzi, M. Dardiry, T. Renahan, D. R. Sharma, R. J. Sommer, Environmental influence on *Pristionchus pacificus* mouth form through different culture methods. Sci. Rep. 7, 1–12 (2017).

46. M. Lenuzzi, H. Witte, M. Riebesell, C. Rödelsperger, R. L. Hong, R. J. Sommer, Influence of environmental temperature on mouth-form plasticity in *Pristionchus pacificus* acts through daf-11-dependent cGMP signaling. J. Exp. Zool. Part B Mol. Dev. Evol. 340, 214–224 (2023).

47. N. Bose, A. Ogawa, S. H. Von Reuss, J. J. Yim, E. J. Ragsdale, R. J. Sommer, F. C. Schroeder, Complex small-molecule architectures regulate phenotypic plasticity in a nematode. Angew. Chemie - Int. Ed. 51, 12438–12443 (2012).

48. M. Dardiry, V. Piskobulu, A. Kalirad, R. J. Sommer, Experimental and theoretical support for costs of plasticity and phenotype in a nematode cannibalistic trait. Evol. Lett. 7, 48–57 (2023).

49. N. Akduman, J. W. Lightfoot, W. Röseler, H. Witte, W. S. Lo, C. Rödelsperger, R. J. Sommer, Bacterial vitamin B12 production enhances nematode predatory behavior. ISME J. 14, 1494–1507 (2020).

50. T. Beltran, V. Shahrezaei, V. Katju, P. Sarkies, Epimutations driven by small RNAs arise frequently but most have limited duration in *Caenorhabditis elegans*. *Nat*. Ecol. Evol. 4, 1539–1548 (2020).

51. A. M. Weller, C. Rödelsperger, G. Eberhardt, R. I. Molnar, R. J. Sommer, Opposing forces of A/T-biased mutations and G/C-biased gene conversions shape the genome of the nematode *Pristionchus pacificus*. Genetics 196, 1145–1152 (2014).

52. R. Kaletsky, R. S. Moore, G. D. Vrla, L. R. Parsons, Z. Gitai, C. T. Murphy, *C. elegans* interprets bacterial non-coding RNAs to learn pathogenic avoidance. Nature 586, 445–451 (2020).

53. O. Rechavi, L. Houri-Ze’Evi, S. Anava, W. S. S. Goh, S. Y. Kerk, G. J. Hannon, O. Hobert, Starvation-induced transgenerational inheritance of small RNAs in *C. elegans*. Cell 158, 277–287 (2014).

54. E. Moreno, B. Sieriebriennikov, H. Witte, C. Rödelsperger, J. W. Lightfoot, R. J. Sommer, Regulation of hyperoxia-induced social behaviour in *Pristionchus pacificus* nematodes requires a novel cilia-mediated environmental input. Sci. Rep. 7, 1–13 (2017).

55. C. Y. Shi, E. R. Kingston, B. Kleaveland, D. H. Lin, M. W. Stubna, D. P. Bartel, The ZSWIM8 ubiquitin ligase mediates target-directed microRNA degradation. Science 370, 1–10 (2020).

56. J. Han, C. A. Lavigne, B. T. Jones, H. Zhang, F. Gillett, J. T. Mendell, A ubiquitin ligase mediates target-directed microRNA decay independently of tailing and trimming. Science 370 (2020).

57. E. R. Kingston, L. W. Blodgett, D. P. Bartel, Endogenous transcripts direct microRNA degradation in *Drosophila*, and this targeted degradation is required for proper embryonic development. Mol. Cell 82, 3872–3884 (2022).

58. Z. Wang, Y. Hou, X. Guo, M. vanderVoet, M. Boxem, J. E. Dixon, A. D. Chisholm, Y. Jin, The EBAX-type Cullin-RING E3 Ligase and Hsp90 Guard the Protein Quality of the SAX-3/Robo Receptor in Developing Neurons. Neuron 79, 903–916 (2013).

59. B. Sieriebriennikov, G. V. Markov, H. Witte, R. J. Sommer, The Role of DAF-21/Hsp90 in Mouth-Form Plasticity in *Pristionchus pacificus*. Mol. Biol. Evol. 34, 1644–1653 (2017).

60. C. Y. Shi, L. E. Elcavage, R. R. Chivukula, J. Stefano, B. Kleaveland, D. P. Bartel, ZSWIM8 destabilizes many murine microRNAs and is required for proper embryonic growth and development. Genome Res. 33, 1482–1496 (2023).

61. P. H. Wu, P. D. Zamore, To Degrade a MicroRNA, Destroy Its Argonaute Protein. Mol. Cell 81, 223–225 (2021).

62. J. Han, J. T. Mendell, MicroRNA turnover: a tale of tailing, trimming, and targets. Trends Biochem. Sci. 48, 26–39 (2023).

63. S. Iwasaki, M. Kobayashi, M. Yoda, Y. Sakaguchi, S. Katsuma, T. Suzuki, Y. Tomari, Hsc70/Hsp90 chaperone machinery mediates ATP-dependent RISC loading of small RNA duplexes. Mol. Cell 39, 292–299 (2010).

64. X. Liu, Y. Y. Yang, Y. Wang, HSP90 and Aha1 modulate microRNA maturation through promoting the folding of Dicer1. Nucleic Acids Res. 50, 6990–7001 (2022).

65. R. Ahmed, Z. Chang, A. E. Younis, C. Langnick, N. Li, W. Chen, N. Brattig, C. Dieterich, Conserved miRNAs are candidate post-transcriptional regulators of developmental arrest in free-living and parasitic nematodes. Genome Biol. Evol. 5, 1246–1260 (2013).

66. E. De Wit, S. E. V. Linsen, E. Cuppen, E. Berezikov, Repertoire and evolution of miRNA genes in four divergent nematode species. Genome Res. 19, 2064–2074 (2009).

67. A. Kozomara, M. Birgaoanu, S. Griffiths-Jones, MiRBase: From microRNA sequences to function. Nucleic Acids Res. 47, D155–D162 (2019).

68. A. Kozomara, S. Griffiths-Jones, MiRBase: Annotating high confidence microRNAs using deep sequencing data. Nucleic Acids Res. 42, 68–73 (2014).

69. S. Griffiths-Jones, R. J. Grocock, S. van Dongen, A. Bateman, A. J. Enright, miRBase: microRNA sequences, targets and gene nomenclature. Nucleic Acids Res. 34, 140–144 (2006).

70. S. Griffiths-Jones, H. K. Saini, S. Van Dongen, A. J. Enright, miRBase: Tools for microRNA genomics. Nucleic Acids Res. 36, 154–158 (2008).

71. B. F. Donnelly, B. Yang, A. L. Grimme, K. F. Vieux, C. Y. Liu, L. Zhou, K. McJunkin, The developmentally timed decay of an essential microRNA family is seed-sequence dependent. Cell Rep. 40 (2022).

72. E. Alvarez-Saavedra, H. R. Horvitz, Many Families of C. elegans MicroRNAs Are Not Essential for Development or Viability. Curr. Biol. 20, 367–373 (2010).

73. M. Liu, P. Liu, L. Zhang, Q. Cai, G. Gao, W. Zhang, Z. Zhu, D. Liu, Q. Fan, *Mir-35* is involved in intestine cell G1/S transition and germ cell proliferation in *C. elegans*. Cell Res. 21, 1605–1618 (2011).

74. D. R. Sharma, W. Roeseler, H. Witte, M. S. Werner, R. J. Sommer, Developmental small RNA transcriptomics reveals divergent evolution of the conserved microRNA *miR-100* and the *let-7*-complex in nematodes. bioRxiv, doi: 10.1101/2024.07.19.604269 [Preprint] (2024).

75. B. Kleaveland, C. Y. Shi, J. Stefano, D. P. Bartel, A Network of Noncoding Regulatory RNAs Acts in the Mammalian Brain. Cell 174, 350–362 (2018).

76. C. T. Neilsen, G. J. Goodall, C. P. Bracken, “IsomiRs - The overlooked repertoire in the dynamic microRNAome” (2012); 10.1016/j.tig.2012.07.005.

77. P. Y. Wang, D. P. Bartel, A statistical approach for identifying primary substrates of ZSWIM8-mediated microRNA degradation in small-RNA sequencing data. BMC Bioinformatics 24, 1–18 (2023).

78. A. Bitetti, A. C. Mallory, E. Golini, C. Carrieri, H. Carreño Gutiérrez, E. Perlas, Y. A. Pérez-Rico, G. P. Tocchini-Valentini, A. J. Enright, W. H. J. Norton, S. Mandillo, D. O’Carroll, A. Shkumatava, MicroRNA degradation by a conserved target RNA regulates animal behavior. Nat. Struct. Mol. Biol. 25, 244–251 (2018).

79. F. Ghini, C. Rubolino, M. Climent, I. Simeone, M. J. Marzi, F. Nicassio, Endogenous transcripts control miRNA levels and activity in mammalian cells by target-directed miRNA degradation. Nat. Commun. 9 (2018).

80. L. Li, P. Sheng, T. Li, C. J. Fields, N. M. Hiers, Y. Wang, J. Li, C. M. Guardia, J. D. Licht, M. Xie, Widespread microRNA degradation elements in target mRNAs can assist the encoded proteins. Genes Dev. 35, 1595–1609 (2021).

81. Z. Han, W. S. Lo, J. W. Lightfoot, H. Witte, S. Sun, R. J. Sommer, Improving transgenesis efficiency and CRISPR-associated tools through codon optimization and native intron addition in *Pristionchus* nematodes. Genetics 216, 947–956 (2020).

82. K. Kagias, R. Pocock, MicroRNA regulation of the embryonic hypoxic response in *Caenorhabditis elegans*. Sci. Rep. 5, 1–9 (2015).

83. V. Susoy, E. J. Ragsdale, N. Kanzaki, R. J. Sommer, Rapid diversification associated with a macroevolutionary pulse of developmental plasticity. Elife 4, 7–17 (2015).

84. V. Susoy, M. Herrmann, N. Kanzaki, M. Kruger, C. N. Nguyen, C. Rödelsperger, W. Röseler, C. Weiler, R. M. Giblin-Davis, E. J. Ragsdale, R. J. Sommer, Large-scale diversification without genetic isolation in nematode symbionts of figs. Sci. Adv. 2, 1–11 (2016).

85. S. Wighard, H. Witte, R. J. Sommer, Conserved switch genes that arose via whole-genome duplication regulate a cannibalistic nematode morph. Sci. Adv. 10, 1–9 (2024).

86. A. Holz, A. Streit, Gain and loss of small RNA classes-characterization of small RNAs in the parasitic nematode family Strongyloididae. Genome Biol. Evol. 9, 2826–2843 (2017).

87. A. Grishok, H. Tabara, C. C. Mello, Genetic requirements for inheritance of RNAi in *C. elegans*. Science 287, 2494–2497 (2000).

88. S. Devanapally, P. Raman, M. Chey, S. Allgood, F. Ettefa, M. Diop, Y. Lin, Y. E. Cho, A. M. Jose, Mating can initiate stable RNA silencing that overcomes epigenetic recovery. Nat. Commun. 12, 1–16 (2021).

89. J. K. Kruschke, Doing Bayesian Data Analysis: A Tutorial with R, JAGS, and Stan, Second Edition (Elsevier Science, 2014).

90. O. Abril-Pla, V. Andreani, C. Carroll, L. Dong, C. J. Fonnesbeck, M. Kochurov, R. Kumar, J. Lao, C. C. Luhmann, O. A. Martin, M. Osthege, R. Vieira, T. Wiecki, R. Zinkov, PyMC: a modern, and comprehensive probabilistic programming framework in Python. PeerJ Comput. Sci. 9, 1–35 (2023).

91. C. R. Harris, K. J. Millman, S. J. van der Walt, R. Gommers, P. Virtanen, D. Cournapeau, E. Wieser, J. Taylor, S. Berg, N. J. Smith, R. Kern, M. Picus, S. Hoyer, M. H. van Kerkwijk, M. Brett, A. Haldane, J. F. del Río, M. Wiebe, P. Peterson, P. Gérard-Marchant, K. Sheppard, T. Reddy, W. Weckesser, H. Abbasi, C. Gohlke, T. E. Oliphant, Array programming with NumPy. Nature 585, 357–362 (2020).

92. J. K. Kruschke, Bayesian Analysis Reporting Guidelines. *Nat*. Hum. Behav. 5, 1282–1291 (2021).

93. H. Li, R. Durbin, Fast and accurate short read alignment with Burrows-Wheeler transform. Bioinformatics 25, 1754–1760 (2009).

94. R. Poplin, V. Ruano-Rubio, M. A. DePristo, T. J. Fennell, M. O. Carneiro, G. A. Van der Auwera, D. E. Kling, L. D. Gauthier, A. Levy-Moonshine, D. Roazen, K. Shakir, J. Thibault, S. Chandran, C. Whelan, M. Lek, S. Gabriel, M. J. Daly, B. Neale, D. G. MacArthur, E. Banks, Scaling accurate genetic variant discovery to tens of thousands of samples. bioRxiv, doi: 10.1101/201178 [Preprint] (2017).

95. P. Danecek, J. K. Bonfield, J. Liddle, J. Marshall, V. Ohan, M. O. Pollard, A. Whitwham, T. Keane, S. A. McCarthy, R. M. Davies, H. Li, Twelve years of SAMtools and BCFtools. Gigascience 10, 1–4 (2021).

96. A. Pires da Silva, “*Pristionchus pacificus* protocols” in WormBook (2013; http://www.wormbook.org/chapters/www_ppageneticprotocols.2/ppageneticprotocols.htm l), pp. 1–20.

97. H. Li, B. Handsaker, A. Wysoker, T. Fennell, J. Ruan, N. Homer, G. Marth, G. Abecasis, R. Durbin, The Sequence Alignment/Map format and SAMtools. Bioinformatics 25, 2078–2079 (2009).

98. R. Rae, H. Witte, C. Rödelsperger, R. J. Sommer, The importance of being regular: *Caenorhabditis elegans* and *Pristionchus pacificus* defecation mutants are hypersusceptible to bacterial pathogens. Int. J. Parasitol. 42, 747–753 (2012).

99. H. Witte, E. Moreno, C. Rödelsperger, J. Kim, J. S. Kim, A. Streit, R. J. Sommer, Gene inactivation using the CRISPR/Cas9 system in the nematode *Pristionchus pacificus*. Dev. Genes Evol. 225, 55–62 (2014).

100. F. Gabler, S. Nam, S. Till, M. Mirdita, M. Steinegger, J. Söding, A. N. Lupas, V. Alva, Protein Sequence Analysis Using the MPI Bioinformatics Toolkit. Curr. Protoc. Bioinforma. 72, 1–30 (2020).

101. J. Trifinopoulos, L. T. Nguyen, A. von Haeseler, B. Q. Minh, W-IQ-TREE: a fast online phylogenetic tool for maximum likelihood analysis. Nucleic Acids Res. 44, W232–W235 (2016).

102. L. T. Nguyen, H. A. Schmidt, A. Von Haeseler, B. Q. Minh, IQ-TREE: A fast and effective stochastic algorithm for estimating maximum-likelihood phylogenies. Mol. Biol. Evol. 32, 268–274 (2015).

103. G. E. Janssens, X. X. Lin, L. Millan-Ariño, A. Kavšek, I. Sen, R. I. Seinstra, N. Stroustrup, E. A. A. Nollen, C. G. Riedel, Transcriptomics-Based Screening Identifies Pharmacological Inhibition of Hsp90 as a Means to Defer Aging. Cell Rep. 27, 467–480.e6 (2019).

104. D. Kim, B. Langmead, S. L. Salzberg, HISAT: A fast spliced aligner with low memory requirements. Nat. Methods 12, 357–360 (2015).

105. Y. Liao, G. K. Smyth, W. Shi, FeatureCounts: An efficient general purpose program for assigning sequence reads to genomic features. Bioinformatics 30, 923–930 (2014).

106. M. I. Love, W. Huber, S. Anders, Moderated estimation of fold change and dispersion for RNA-seq data with DESeq2. Genome Biol. 15, 1–21 (2014).

107. M. Marcel, Cutadapt removes adapter sequences from high-throughput sequencing reads. EMBnet J. 17, 10–12 (2011).

108. B. Langmead, C. Wilks, V. Antonescu, R. Charles, Scaling read aligners to hundreds of threads on general-purpose processors. Bioinformatics 35, 421–432 (2019).

109. A. Kozomara, S. Griffiths-Jones, MiRBase: Integrating microRNA annotation and deep-sequencing data. Nucleic Acids Res. 39, 152–157 (2011).

110. W. Li, A. Godzik, Cd-hit: A fast program for clustering and comparing large sets of protein or nucleotide sequences. Bioinformatics 22, 1658–1659 (2006).

111. C. Rödelsperger, J. M. Meyer, N. Prabh, C. Lanz, F. Bemm, R. J. Sommer, Single-Molecule Sequencing Reveals the Chromosome-Scale Genomic Architecture of the Nematode Model Organism *Pristionchus pacificus*. Cell Rep. 21, 834–844 (2017).

112. S. Koren, B. P. Walenz, K. Berlin, J. R. Miller, N. H. Bergman, A. M. Phillippy, Canu: Scalable and accurate long-read assembly via adaptive κ-mer weighting and repeat separation. Genome Res. 27, 722–736 (2017).

113. M. Dardiry, G. Eberhard, H. Witte, C. Rödelsperger, J. W. Lightfoot, R. J. Sommer, Divergent combinations of cis-regulatory elements control the evolution of phenotypic plasticity. PLoS Biol. 21, 1–11 (2023).

114. A. McGaughran, C. Rödelsperger, D. G. Grimm, J. M. Meyer, E. Moreno, K. Morgan, M. Leaver, V. Serobyan, B. Rakitsch, K. M. Borgwardt, R. J. Sommer, Genomic Profiles of Diversification and Genotype-Phenotype Association in Island Nematode Lineages. Mol. Biol. Evol. 33, 2257–2272 (2016).

115. C. Rödelsperger, The community-curated *Pristionchus pacificus* genome facilitates automated gene annotation improvement in related nematodes. BMC Genomics 22, 1–12 (2021).

116. M. Athanasouli, H. Witte, C. Weiler, T. Loschko, G. Eberhardt, R. J. Sommer, C. Rödelsperger, Comparative genomics and community curation further improve gene annotations in the nematode *Pristionchus pacificus*. BMC Genomics 21, 1–9 (2020).

117. N. Prabh, C. Rödelsperger, Multiple *Pristionchus pacificus* genomes reveal distinct evolutionary dynamics between de novo candidates and duplicated genes. Genome Res. 32, 1315–1327 (2022).

118. F. A. Simão, R. M. Waterhouse, P. Ioannidis, E. V. Kriventseva, E. M. Zdobnov, BUSCO: Assessing genome assembly and annotation completeness with single-copy orthologs. Bioinformatics 31, 3210–3212 (2015).

119. S. C. Potter, A. Luciani, S. R. Eddy, Y. Park, R. Lopez, R. D. Finn, HMMER web server: 2018 update. Nucleic Acids Res. 46, W200–W204 (2018).

