## Supplemental Figures and Tables for "EBAX-1/ZSWIM8 destabilizes miRNAs resulting in transgenerational memory of a predatory trait"

**The PDF file includes:**

Figs. S1 to S5
Tables S1 to S2

**Other Supplementary Materials for this manuscript include the following:**

Data S1 to S3

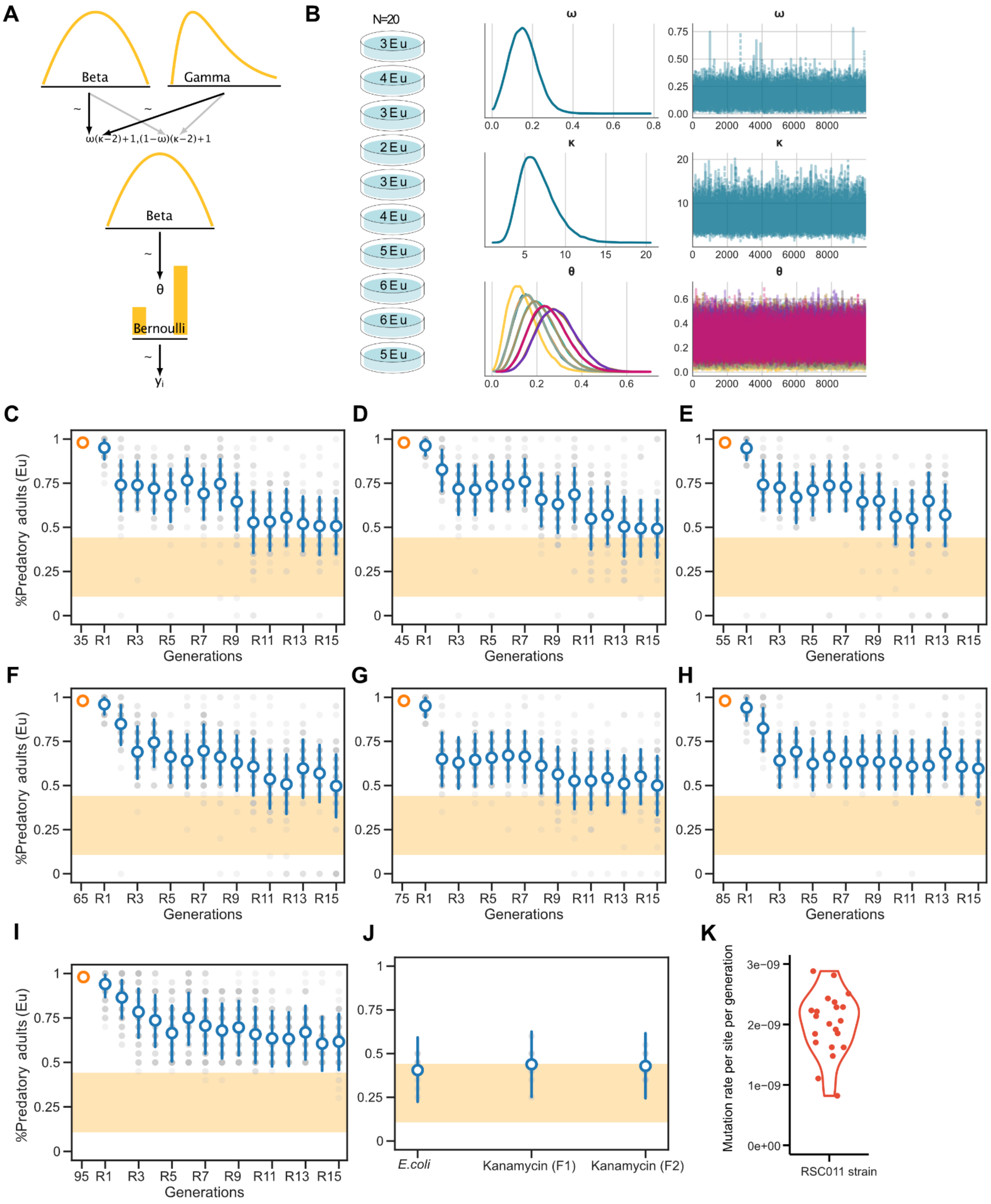
A
Beta
Gamma
ω(κ−2)+1,(1−ω)(κ−2)+1
Beta
θ
Bernoulli
yᵢ
B
N=20
3 Eu
4 Eu
3 Eu
2 Eu
3 Eu
4 Eu
5 Eu
6 Eu
6 Eu
5 Eu
ω
κ
θ
C
D
E
F
G
H
I
J
K
%Predatory adults (Eu)
Generations
35 R1 R3 R5 R7 R9 R11 R13 R15
45 R1 R3 R5 R7 R9 R11 R13 R15
55 R1 R3 R5 R7 R9 R11 R13 R15
65 R1 R3 R5 R7 R9 R11 R13 R15
75 R1 R3 R5 R7 R9 R11 R13 R15
85 R1 R3 R5 R7 R9 R11 R13 R15
95 R1 R3 R5 R7 R9 R11 R13 R15
E.coli
Kanamycin (F1)
Kanamycin (F2)
Mutation rate per site per generation
3e−09
2e−09
1e−09
0e+00
RSC011 strain

**Fig. S1. Diet reversal experiments and mutation rate of RSC011 after 100 generations of single worm transfer.**

(A) The Hierarchical Bayesian model used to infer mean probability estimates of the predatory mouth form. Graphical representation of the model to infer the mean highest density interval (HDI). (B) Plates represent independent replicates of the actual Eu counts. The probability of developing the Eu mouth form ($\theta$) for each replicate was derived by linking all replicates using the hyperparameters, $\omega$ and $\kappa$. The prior distribution on the given parameters is described as $\omega$~Beta ($\alpha$,$\beta$) and $\kappa$~Gamma (3,1). The stability of the estimated $\theta$ values for a given generation was validated using common convergence diagnostics (see Materials and Methods for details). (C-I) Mean probability of the predatory mouth form during food reversal experiments back to *E. coli* after original exposure to *Novosphingobium* for 35, 45, 55, 65, 75, 85 and 95 generations. (J) Mean probability of the predatory mouth form after exposure to kanamycin-resistant *E. coli* strain for two generations. The 95% HDI for the means was estimated using a Bayesian hierarchical model and indicates the probability of expressing the Eu morph inferred from the experimental data. The yellow region displays the upper and lower limit of the baseline Eu response of RSC011 ($0.115 \leq \text{HDI}(\bar{\theta}_{\text{Control}}) \leq 0.436$). (K) Mutation rate per site per generation for *P. pacificus* RSC011. Each dot represents the mutation rate for independent lines sequenced after 100 generations.

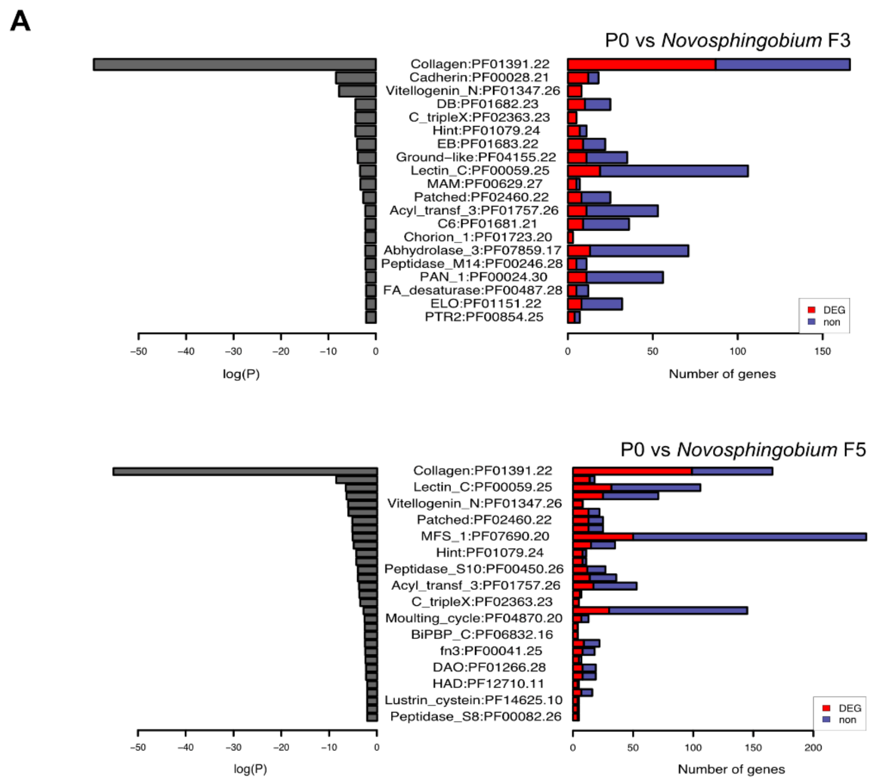


**Fig. S2. Gene enrichment analysis upon *Novosphingobium* exposure.**

(A) Gene-enrichment analysis on *Novosphingobium* for three and five generations. The complete gene lists are listed in Data S1.

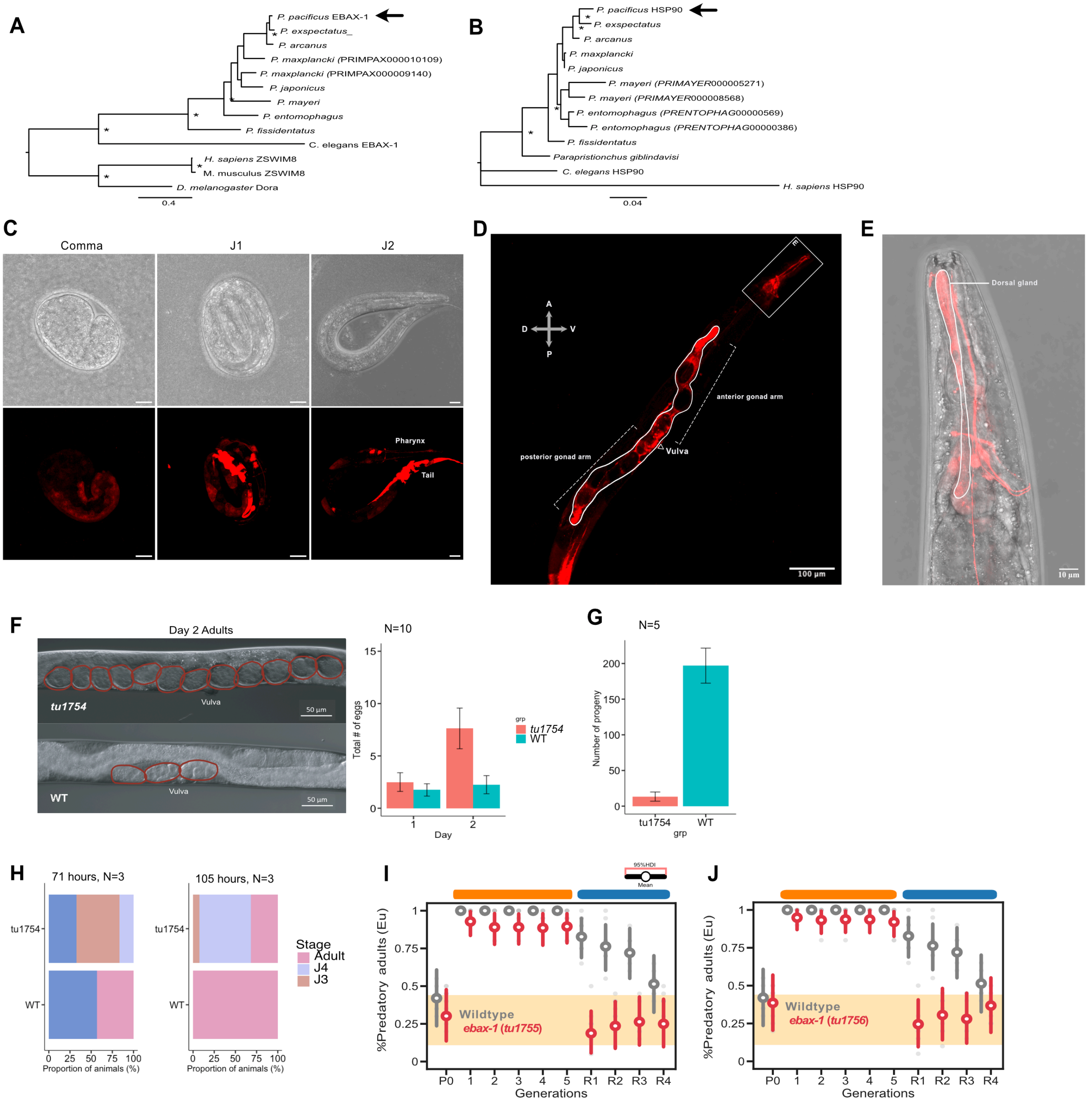


**Fig. S3. Pairwise orthology relationships of the candidate genes and involvement of *ebax-1* in transgenerational inheritance of the predatory morph.**

(A) One-to-one orthology relationships for EBAX-1 and (B) DAF-21. Both maximum likelihood phylogenies are based on the genus *Pristionchus (P. pacificus, P. exspectatus, P. arcanus, P. maxplancki, P. japonicus, P. mayeri, P.entomophagus, P. fissidentatus)* (some includes its 'basal' relative *Parapristionchus*), *C. elegans*, mammals, and insects. Nodes with bootstrap support values of ≥ 90 are labelled with asterisks (*) and arrows indicate *P. pacificus* genes targeted for functional analyses. *P. maxplancki* has two proteins encoding a zinc-finger SWIM-type profile

(PRIMPAX000009140) and a ZSWIM4-8- C-terminal (PRIMPAX000010109), both are included in the EBAX-1 phylogeny. The top two protein BLAST hits of *P. mayeri* and *P. entomophagus* species against *C. elegans* DAF-21 are also included. (C-E) Transcriptional reporter construct of *Ppa-ebax-1*. (C) *Ppa-ebax-1*p::TurboRFP during embryonic development, starting from the comma stage, and J2 stage. Upper panels are DIC images and bottom panels are fluorescent images of the same embryos. Scale bars represent 10 µm. (D) RFP expression on the pharyngeal neurons, vulval region and gonad tissue of adults. The double-arrows in a cross indicate anatomical worm orientation: A-anterior, P-posterior, D-dorsal, V-ventral. (E) Zoom-in on the head region depicted in (D), previous image. (F) Egg-laying-defective phenotype in *Ppa-ebax-1*(*tu1754*) (left). Eggs inside the worm body are outlined in red circles. Average number of accumulated eggs on day-2 *Ppa-ebax-1(tu1754)* adults compared to wild type (right). (G) Average number of progeny counts in *Ppa-ebax-1 (tu1754)* and wild type animals. Error bars are represented in standard deviation. Orange, *Ppa-ebax-1(tu1754)*; green, wild type. (H) Developmental staging of *Ppa-ebax-1(tu1754)* and wild type showing the percentage of J3, J4 and young adults after 71 and 105 hours on *E. coli*. (I-J) Mean probability of the predatory morph in other *Ppa-ebax-1* in-frame alleles for 5 generations of *Novopshingobium* exposure and reversal to *E. coli*. Red, *Ppa-ebax-1(tu1754)*; grey, wild type. The 95% highest density interval (HDI) was used to visualize the probability of observing the Eu morph. The yellow region displays the upper and lower limit of the RSC011 baseline Eu response. More details about the molecular lesions are listed in Table S2.

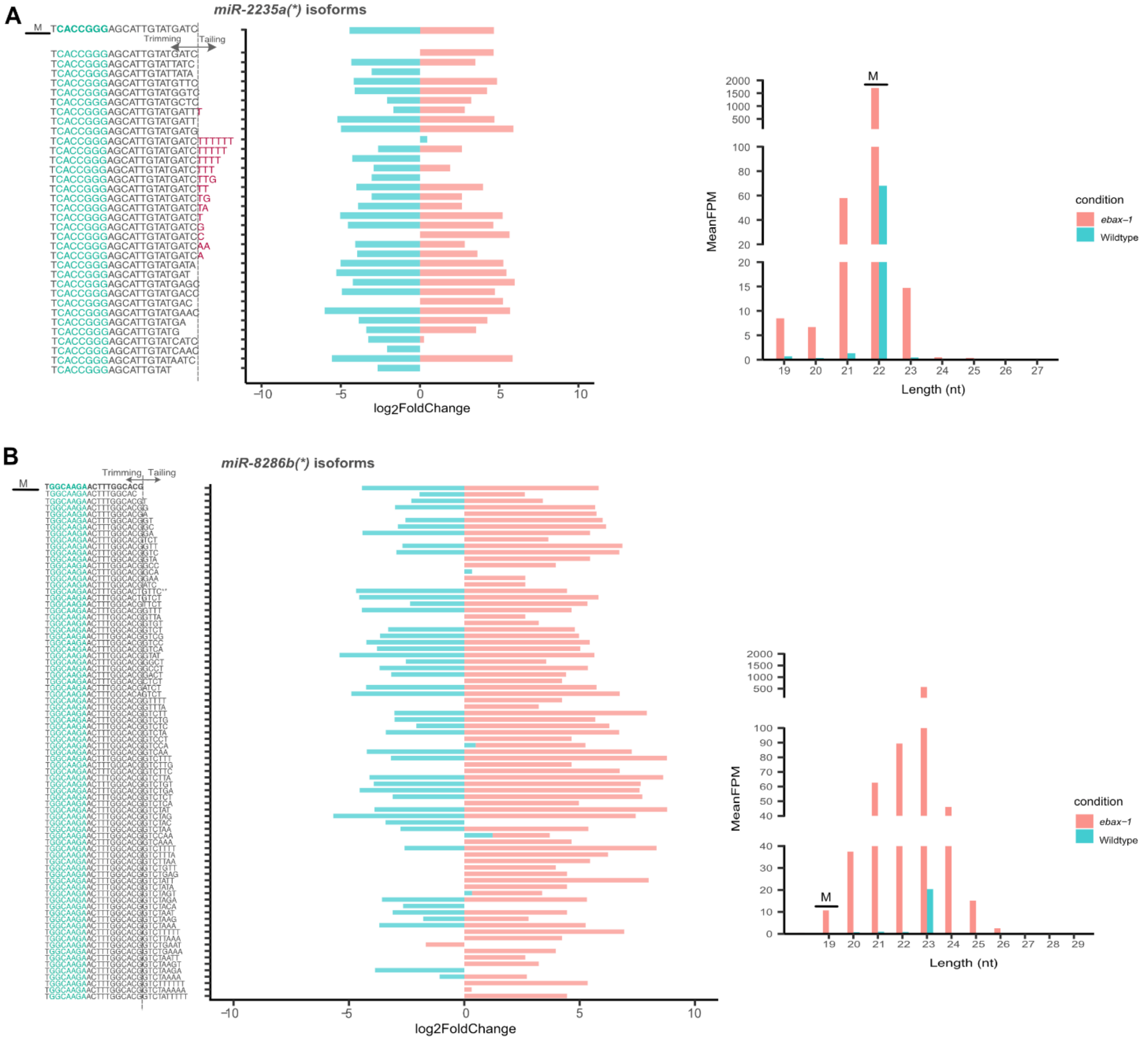


**Figure S4. Influence of *ebax-1* on the distribution and patterns of *miR-8286b* and *miR-2235a* isoforms.**

(A-B) Tailed and trimmed versions of *miR-2235a* and *miR-8286b* during reversal to *E. coli* (left). Plotted are the log2 fold change from wild type (blue) and *ebax-1*(*tu1754*) animals (red). Also plotted are the length distribution and normalized Fragments per Million (FPM) counts of *miR-2235a* and *miR-8286b* reads (left). M represents the designated mature miRNA lengths for each strand are shown.

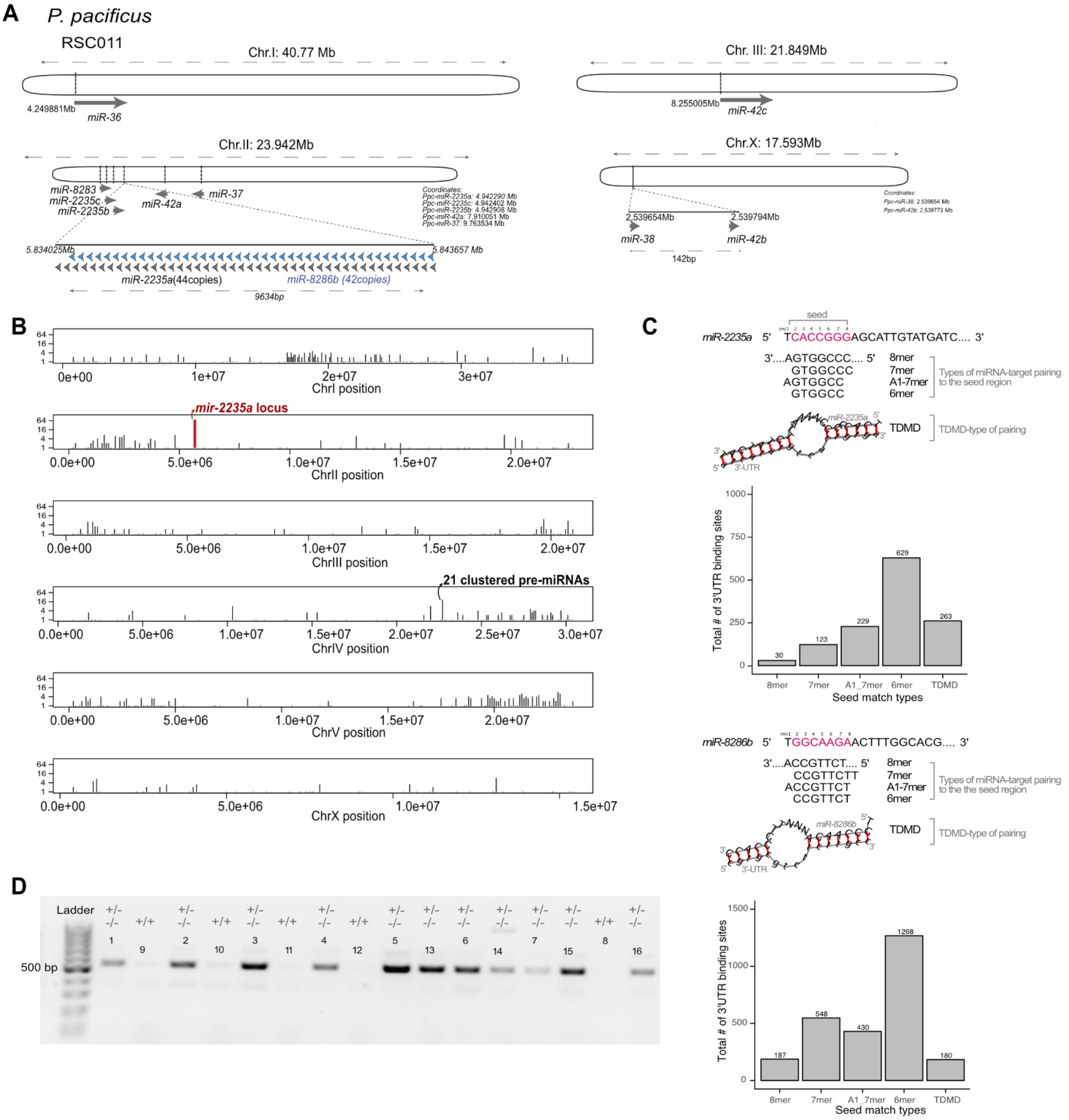


**Figure S5. Earlier and extended predatory mouth form memory of *miR-2235a* deletion alleles**

(A). *miR-35/miR-2235a* family members distributed in different chromosomes. Grey arrows, miR-2235a members. Blue arrows, *miR-8286b*. (B) Graphical representation of precursor miRNA in clusters across the *P. pacificus* genome. *miR-2235a* (red) is identified to be the largest clustered miRNAs in chromosome II, followed by the 21 clustered miRNAs in chromosome IV. (C) 3'-UTR prediction of *miR-2235a* and *miR-8286b* targets. Various types of miRNA target sites are shown with pairings restricted to the seed region and the TDMD-type of pairing with complementarity extending to the 3' region of the miRNA. The complete list of predicted targets are compiled in Data S3. (D) Agarose gel documentation after PCR of progeny individuals from a heterozygous animal for the *miR-2235a* cluster deletion. Lanes without bands are wild type (+/+, without the deletion), with bands of <500 bp are either heterozygous (+/-) or homozygous (-/-) lines for the deletion (see Materials and Methods for details). More details about the molecular lesions are listed in Table S2.

**Table S1. *Ppa-ebax-1* and *Ppa-daf-21* alleles isolated from the genetic screen by EMS mutagenesis.**

| Alelle | Gene ID | *C. elegans* 1:1 ortholog | Position | Reference | Variant | Mutation type | nt substitution |
|---|---|---|---|---|---|---|---|
| *tu803* | RSC011000001067 | *daf-21* | 35424586 | C | A | nonsynonym [E>D] | GAG>GAT |
| *tu797* | RSC011000001067 | *daf-21* | 35425940 | C | T | nonsynonym [G>S] | GGC>AGC |
| *tu770* | RSC011000001067 | *daf-21* | 35425073 | C | T | nonsynonym [E>K] | GAG>AAG |
| *tu776* | RSC011000009259 | *ebax-1* | 7625936 | C | T | nonsynonym [S>F] | TCC>TTC |
| *tu780* | RSC011000009259 | *ebax-1* | 17621261 | C | T | nonsynonym [T>I] | ACT>ATT |
| *tu759* | RSC011000009259 | *ebax-1* | 17626295 | T | A | nonsynonym [V>D] | GTT>GAT |

**Table S2. Molecular lesions of *Ppa-ebax-1* and the *miR-2235a* locus. *tid*, *t*ransgenerational *i*nheritance *d*efective. TEI, transgenerational epigenetic inheritance.**

| Alleles | Gene ID; *C. elegans* best hit | Molecular lesions | Phenotype | sgRNA(PAM) | Forward primer | Reverse primer |
|---|---|---|---|---|---|---|
| *tu1754* | RSC011000009259; *ebax-1* | 10 bp insertion | Egg-laying defective,slow growth, low progeny, *tid* | AAGGAGTAGTAAAAGGATAT (GGG) | GGTGCAAACAAAAGTGACCT | CTGTTGGGATAGACAGCCAG |
| *tu1755* | | 60 bp deletion | *tid* | | | |
| *tu1756* | | T1231I + 12 bp deletion | *tid* | | | |
| *tuDf9* | *miR-2235a* (ChrII:5,834,025-5,843,657) | 9,408 bp deletion | Earlier and extended TEI | GCCCGATCATACAATGCTCC (CGG) | CCCTGCGTACTGACTATCTG | GGTTGGCTGGATAGCGTTTG |
| *tuDf10* | | 9,632 bp deletion | Earlier and extended TEI | | | |
| *tuDf11* | | 9,856 bo deletion | Earlier and extended TEI | | | |

**Data S1. Differential gene expression analysis and enriched gene sets after exposure to *Novosphingobium* for three and five generations.**
(A and B) Upregulated and downregulated gene lists after three and five generations on *Novosphingobium.* (C and D) Lists of Pfam-domains enriched in the differentially expressed gene sets after three and five generations on *Novosphingobium.*

**Data S2. Small RNA sequencing analysis of wildtype and *ebax-1* mutant animals.** (A) List of differentially-expressed sRNAs between P0 on *E. coli* and F10 on *Novosphingobium*. (B) List of differentially-expressed sRNAs between F10 on *Novosphingobium* and two generations on *E. coli* (R2). (C) List of differentially-expressed sRNAs between wild type and *ebax-1* on F10 *Novosphingobium*. (D) List of differentially-expressed sRNAs between wild type and *ebax-1* after reversal to *E. coli*. (E-F) List of *miR-2235a* and *miR-8286a* isoforms during reversal upon *ebax-1* knockout.

**Data S3. Genome-wide 3'-UTR prediction of miRNA targets in RSC011**.
